# Identification of novel wraparound transcripts in JC polyomavirus

**DOI:** 10.1101/2024.10.30.619564

**Authors:** Shun Iida, Kenta Takahashi, Sohtaro Mine, Tadaki Suzuki, Harutaka Katano

## Abstract

JC polyomavirus (JCPyV) is a ubiquitous pathogen causing progressive multifocal leukoencephalopathy (PML) in immunocompromised individuals. Polyomaviruses (PyVs) utilize complex transcriptional regulation mechanisms to express diverse mRNAs from their compact, double-stranded circular DNA genomes. A recent study employed next-generation sequencing (NGS) to provide a detailed transcriptome atlas of PyVs, including BK polyomavirus and simian virus 40. However, detailed information regarding the JCPyV transcriptome remains limited. Here, we conducted a comprehensive analysis combining short-read and long-read NGS methods to unveil the detailed transcriptome atlas of JCPyV. RNA isolated from IMR-32 and 293 cells transfected with the circular JCPyV genome was analyzed by NGS, leading to the discovery of 39 novel viral transcripts in addition to 12 previously annotated ones. Among these novel transcripts, we identified characteristic wraparound transcripts, conserved across PyVs, which are constructed from long primary transcripts generated through continuous, multiple rounds of circular transcription of the viral genome. These wraparound transcripts included both late transcripts harboring leader-to-leader repeated sequences and SuperT transcripts containing multiple LxCxE motifs. Notably, wraparound transcripts, including a SuperT transcript, were also detected in brain tissues from PML patients. Collectively, this study significantly expands our understanding of the JCPyV transcriptome, revealing the expression of wraparound transcripts in PML lesions. These findings provide valuable insights into the molecular basis of JCPyV gene expression and PML pathogenesis, potentially facilitating the development of effective countermeasures against PML.

**Author Summary:** JC polyomavirus (JCPyV) was first identified in 1971 from the brain tissue of a patient with progressive multifocal leukoencephalopathy (PML), a life-threatening neurological disease. Despite over 50 years since the pathogen’s identification, the prognosis for PML remains extremely poor due to the lack of effective therapeutics. Elucidating the JCPyV transcriptome can expand our understanding of viral gene expression and PML pathogenesis, potentially providing targets for novel therapeutic strategies. Here, we employed a combination of three next-generation sequencing technologies to explore novel viral transcripts, creating a detailed transcriptome atlas of JCPyV. We identified 39 novel viral transcripts in addition to the 12 previously annotated ones in cultured cells transfected with the JCPyV genome. These novel transcripts included wraparound transcripts, which arise from continuous, multiple rounds of circular transcription of the viral genome. Wraparound transcripts were characterized by the presence of tandem repeats of approximately 100 nucleotides. Notably, these characteristic wraparound transcripts were also detected in brain tissues from PML patients. Our study provides new insights into JCPyV gene expression, potentially contributing to the development of effective therapeutics.

## Introduction

Polyomavirus (PyV) is a family of viruses with double-stranded circular DNA genomes approximately 5,000 base pairs in size [1]. Among the 13 known human PyVs, at least four, including JC polyomavirus (JCPyV), BK polyomavirus (BKPyV), Merkel cell polyomavirus (MCPyV), and trichodysplasia spinulosa-associated polyomavirus (TSPyV), have been linked to human diseases [2–5]. JCPyV, the causative agent of a life-threatening demyelinating disease progressive multifocal leukoencephalopathy (PML), was first isolated from the brain tissue of a PML patient in 1971 [3]. JCPyV is a ubiquitous pathogen, with approximately 80% of adults in the general population seropositive [6]. However, PML occurs primarily in immunocompromised individuals such as patients of acquired immunodeficiency syndrome (AIDS) [7], and those taking immunomodulatory therapies [8–17].

PyVs utilize intricate transcriptional regulation to encode multiple transcripts from their compact genomes. As with other members of PyV, the JCPyV genome is divided into three distinct regions: the early region, the late region, and the non-coding control region (NCCR) [18]. The early region encodes large T (LT) and small t (ST) antigens, which play crucial roles in viral replication [18–20]. Conversely, the late region is responsible for encoding three viral capsid proteins (VP1, VP2, and VP3) and the agnoprotein (Agno) [18, 21]. Additionally, the JCPyV genome encodes T’135, T’136, and T’165 antigens, which are splicing variants of the early region T antigens, and trans-spliced open reading frames (ORFs) including ORF1 and ORF2 [22, 23]. Expression of the early and the late transcripts are temporarily regulated by the bidirectional promoter located in the NCCR region.

Recent advances in next-generation sequencing (NGS) have elucidated detailed transcriptomic profiles of viruses. Notably, long-read sequencing technologies, such as nanopore sequencing and single-molecule real-time sequencing (SMRTseq), have revealed the complex nature of viral transcriptomes [24]. These findings provide valuable insights into the molecular basis of virus-related diseases and could potentially serve as novel therapeutic targets. In the context of PyVs, a recent study combining short-read and long-read sequencing technologies unraveled the detailed transcriptome atlas of BKPyV and simian virus 40 (SV40), revealing complex patterns of viral transcript expression, including wraparound transcripts such as SuperT [25]. Although the previous report provided insightful observations into transcriptomics of BKPyV and SV40, our understanding of JCPyV transcriptomics remains limited.

In this study, we conducted a comprehensive analysis of the JCPyV transcriptome using both short-read and long-read RNAsequencing technologies, creating a detailed transcript atlas of JCPyV. Additionally, we investigated the expression of newly identified wraparound transcripts, including SuperT, in brain tissues obtained from patients with PML.

## Results

### JCPyV-encoded transcripts are detected by NGS from both IMR-32 and 293 cells

We utilized NGS to analyze RNA samples purified from cells transfected with the circular genomic DNA of the Mad-1 strain of JCPyV [3], aiming to generate a comprehensive transcriptome atlas of the virus (**Fig 1A**). IMR-32 cells, known for their efficient JCPyV replication [26], were used, and the results were subsequently validated in 293 cells. We initially mapped sequence reads obtained from three distinct NGS methods, including short-read RNA sequencing (short-RNAseq), direct RNA sequencing (dRNAseq), and SMRTseq, to the JCPyV genome. Read mapping revealed that long-read sequencing reads (dRNAseq and SMRTseq) aligned clearly to the JCPyV ORFs (**Fig 1B**). SMRTseq exhibited a relatively higher read depth at the 5’ end, possibly due to the difference in read lengths compared to dRNAseq (**S1 Fig**). While short-read sequencing reads largely consistent with long-read sequencing results, it showed more prominent noise (**Fig 1B**). In IMR-32 cells, both poly(A)+ and ribo-depleted RNA short-read sequencing yielded over 100 million reads; however, less than 0.1% spanned a splicing junction and mapped to the JCPyV genome (**S1 Table**). As for SMRTseq, 1.98% (67,848 reads) of the total 3,430,793 reads mapped to viral transcripts, with 0.91% (31,237 reads) representing spliced transcripts (**S1 Table**). Conversely, dRNAseq identified fewer JCPyV transcript-derived reads (3,284 total and 1,028 spliced) (**S1 Table**). These findings were similarly replicated in 293 cells (**S2 Table**). Collectively, these results indicate that RNA extracted from JCPyV genomic DNA-transfected IMR-32 and 293 cells is suitable for investigating viral transcripts.

**Fig 1:**
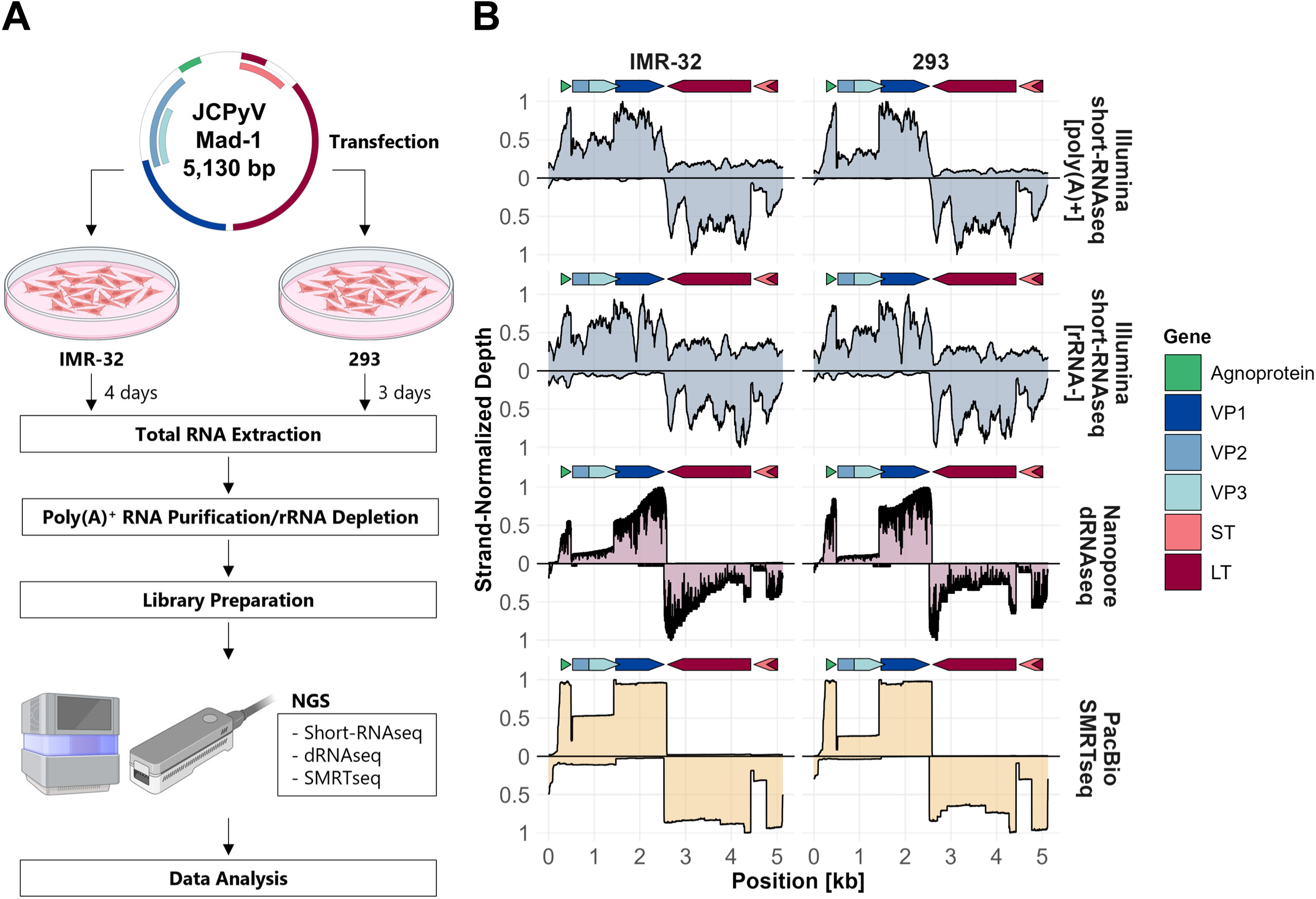
Read mapping to the JCPyV genome. **A.** A schematic overview of this study is shown. Total RNA was extracted from IMR-32 and 293 cells transfected with circular genomic DNA of the JCPyV Mad-1 strain. The RNA underwent poly(A)+ RNA purification or ribosomal RNA (rRNA) depletion, followed by NGS library construction. Sequencing data obtained by short-read RNA sequencing (short-RNAseq), direct RNA sequencing (dRNAseq), and single-molecule real-time sequencing (SMRTseq) were subjected to further analysis. This figure was created with BioRender.com. **B.** Sequence coverage of each method was assessed by mapping the sequence reads from short-RNAseq, dRNAseq, and SMRTseq to the JCPyV genome. Read depth is shown as a relative value, with the highest value on the positive/negative strand normalized to 1. Arrows above each panel indicate the ORFs in the JCPyV genome.

### Identification of novel transcripts encoded by JCPyV

To identify novel JCPyV-encoded transcripts, we further analyzed the NGS data. We found a total of 45 transcripts in IMR-32 cells, including 33 novel and 12 previously annotated ones (**Fig 2A**). Notably, 20 transcripts were detected by both dRNAseq and SMRTseq, with 11 unique to SMRTseq and 2 unique to dRNAseq. A similar trend was observed in 293 cells, where we identified 36 transcripts: 25 novel and 11 previously annotated (**Fig 2B**). In both cell lines, SMRTseq detected more transcripts than dRNAseq, likely due to the relatively low sequencing depth observed at the 5’ end of transcripts in dRNAseq (**Fig 1B**). Combining results from both cell lines, we identified 39 novel viral transcripts, with 14 unique to IMR-32 and 6 unique to 293 cells (**Fig 2C**). We examined transcript abundance across sequencing methods in both cell lines. The cumulative proportion of reads revealed that the 10 most abundant transcripts account for approximately 90% of total reads in all methods, for both early and late strands (**S2 Fig**).

**Fig 2:**
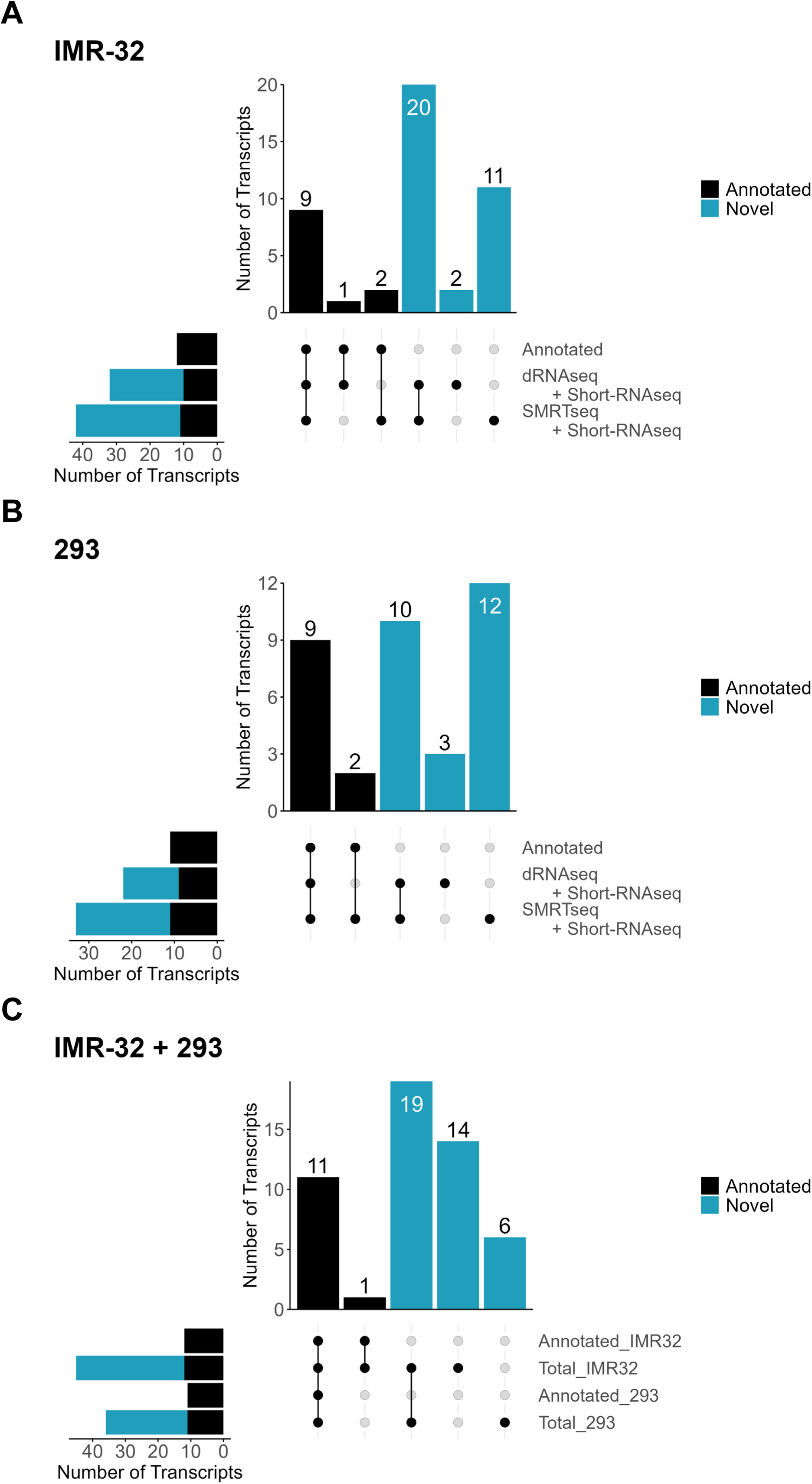
Counts of annotated JCPyV transcripts identified by NGS. **A-B.** Upset plots indicating the numbers of annotated transcripts, direct RNA sequencing (dRNAseq)-detected transcripts, and single-molecule real-time sequencing (SMRTseq)-detected transcripts. The results from IMR-32 cells **(A)** and 293 cells **(B)** are presented. **C.** Upset plot indicating intersections between annotated and novel transcripts, and the cells from which the transcripts were detected. Short-RNAseq, short-read RNA sequencing.

Next, we summarized the detailed profile of viral transcripts detected in this study (**Table 1**) and generated a transcriptome atlas (**Fig 3A, S3A, S4A, S5A, and S6A Figs**). The relative abundance of each transcript is also summarized (**Fig 3B, S3B, S4B, S5B, and S6B Figs**). The novel transcripts from the early region of the JCPyV genome, which lack a large portion of the LT exon, exhibited similar structures, whereas the novel transcripts from the late region demonstrated heterogeneity in their structure (**Fig 3A**). This diversity among late transcripts partly depends on novel splice sites located at positions 298, 376, and 400. Interestingly, the splice site at position 400 served as both a splicing donor (for IMR-L21) and a splicing acceptor (for IMR-L9, IMR-L13, IMR-L18, and VP1-401 wraparound transcripts) (**Fig 3A**). Notably, some of the novel transcripts contained wraparound transcripts harboring a tandem repeat of approximately 100 nucleotides (nt) (**Fig 3A**). Some of the novel transcripts from the late region are expressed more abundantly than several previously known transcripts, such as M5 and M6, and the majority of these are wraparound transcripts (**Fig 3B**). In contrast, novel transcripts from the early region were less abundant than all the known transcripts (**Fig 3B**). Taken together, these data provide a transcriptome atlas of JCPyV, revealing a far greater complexity of viral transcripts than previously recognized.

**Fig 3:**
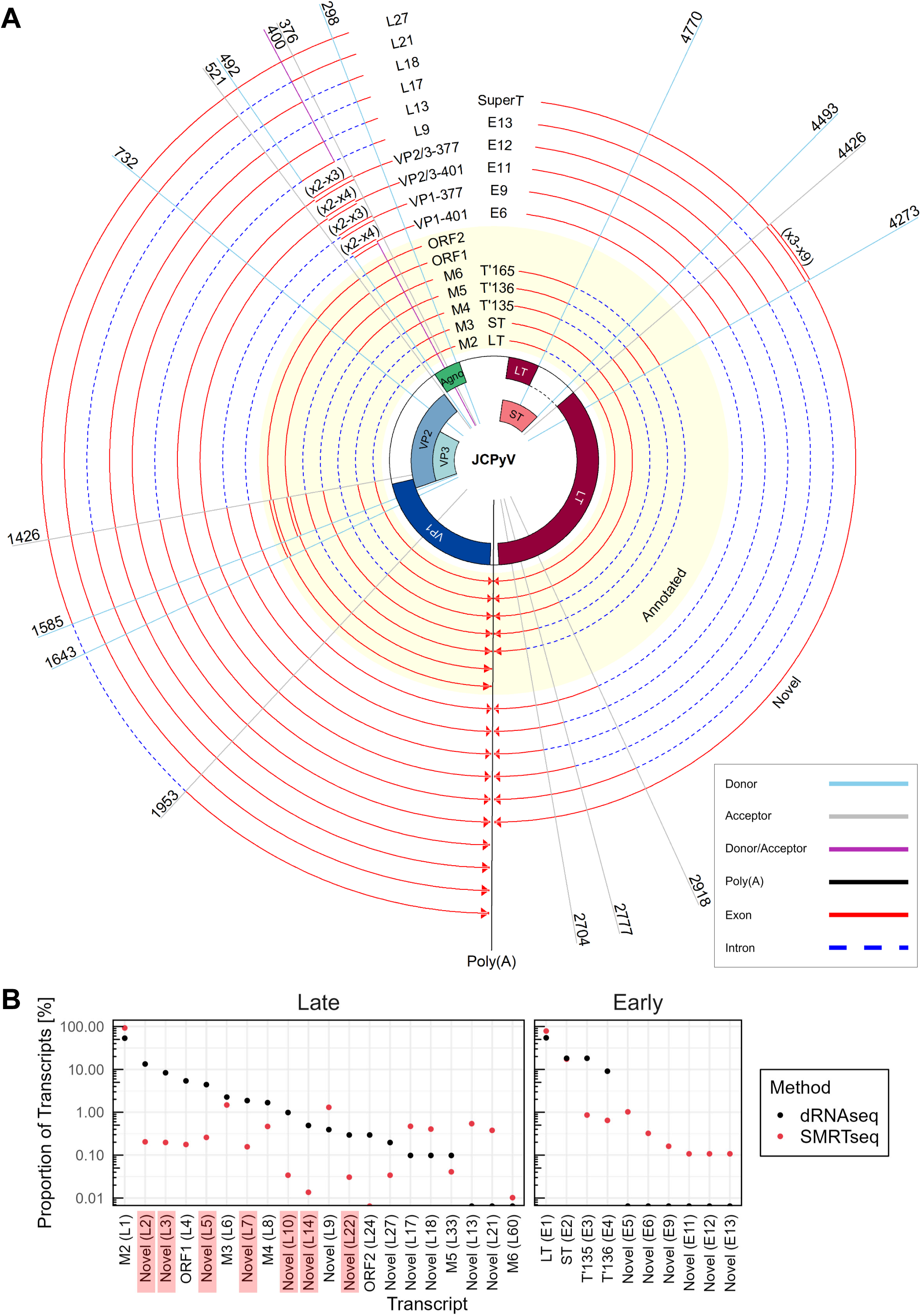
Overview of the JCPyV transcripts. **A.** The JCPyV transcripts detected in this study are summarized in this panel. Each arrow represents a viral transcript, with red regions indicating exons and blue dashed regions indicating introns, respectively. Repeating units of wraparound transcripts are shown as double lines. The transcripts highlighted with yellow background were annotated previously, while others are novel. Radial lines indicate splicing donors/acceptors, or the approximate poly(A) sites. ORFs in the JCPyV genome are shown at the center. **B.** Relative abundance of viral transcripts, demonstrated in **Fig 3A**, in IMR-32 cells. The IDs of wraparound transcripts are highlighted with red background. dRNAseq, direct RNA sequencing; SMRTseq, single-molecule real-time sequencing.

**Table 1:** JCPyV-encoded transcripts detected in this study.

| ID (IMR-32) | ID (293) | Description | Early/Late | Introns |
| --- | --- | --- | --- | --- |
| IMR-E1 | 293-E1 | LT | Early | 4770_4426 |
| IMR-E2 | 293-E2 | ST | Early | 4493_4426 |
| IMR-E3 | 293-E4 | T'135 | Early | 4273_2918<br>4770_4426 |
| IMR-E4 | 293-E3 | T'136 | Early | 4273_2777<br>4770_4426 |
| IMR-E5 | 293-E5 | T'165 | Early | 4273_2704<br>4770_4426 |
| IMR-E6 | 293-E11 | Novel | Early | 4273_2918 |
| IMR-E7 | NA | Novel | Early | 1420_484 |
| IMR-E8 | NA | Novel | Early | 5294_4426 |
| IMR-E9 | 293-E7 | Novel | Early | 4273_2777<br>4493_4426 |
| IMR-E10 | NA | Novel | Early | 9403_4426<br>9900_9556 |
| IMR-E11 | 293-E12 | Novel | Early | 4273_2704<br>4493_4426 |
| IMR-E12 | 293-E8 | Novel | Early | 4273_2777 |
| IMR-E13 | 293-E10 | Novel | Early | 4273_2918<br>4493_4426 |
| NA | 293-E6 | Novel | Early | 4273_2704 |
| NA | 293-E9 | Novel | Early | 4273_2798<br>4770_4426 |
| NA | 293-E13 | Novel | Early | 4273_2817<br>4770_4426 |
| NA | 293-E14 | Novel | Early | 4493_4426<br>4770_4585 |
| NA | 293-E15 | Novel | Early | 4273_2817<br>4493_4426 |
| IMR-L1 | 293-L1 | M2 | Late | 492_1426 |
| IMR-L2 | 293-L3 | Novel | Late | 492_5530<br>5622_6556 |
| IMR-L3 | 293-L2 | Novel | Late | 492_5506<br>5622_6556 |
| IMR-L4 | 293-L5 | ORF1 | Late | 492_1426<br>1585_5651 |
| IMR-L5 | 293-L9 | Novel | Late | 492_5530 |
| IMR-L6 | 293-L4 | M3 | Late | 492_1426<br>1585_1953 |
| IMR-L7 | 293-L10 | Novel | Late | 492_5506 |
| IMR-L8 | 293-L6 | M4 | Late | 492_1953 |
| IMR-L9 | 293-L8 | Novel | Late | 298_400<br>492_1426 |
| IMR-L10 | 293-L16 | Novel | Late | 492_5530<br>5622_10660<br>10752_11686 |
| IMR-L11 | NA | Novel | Late | 1585_5651 |
| IMR-L12 | NA | Novel | Late | 492_5530<br>5622_10636<br>10752_11686 |
| IMR-L13 | 293-L19 | Novel | Late | 298_400 |
| IMR-L14 | 293-L7 | Novel | Late | 492_5506<br>5622_10636<br>10752_11686 |
| IMR-L15 | NA | Novel | Late | 492_5530<br>5622_6556<br>6715_10781 |
| IMR-L16 | NA | Novel | Late | 492_5530<br>5622_7083 |
| IMR-L17 | 293-L17 | Novel | Late | 298_521 |
| IMR-L18 | 293-L11 | Novel | Late | 315_400<br>492_1426 |
| IMR-L19 | NA | Novel | Late | 1643_5651 |
| IMR-L20 | NA | Novel | Late | 492_5506<br>5622_6556<br>6715_7083 |
| IMR-L21 | 293-L18 | Novel | Late | 400_521 |
| IMR-L22 | 293-L14 | Novel | Late | 492_5530<br>5622_10660 |
| IMR-L23 | NA | Novel | Late | 492_5506<br>5622_10660<br>10752_11686 |
| IMR-L24 | NA | ORF2 | Late | 492_1426<br>1643_5651 |
| IMR-L25 | NA | Novel | Late | 492_5530<br>5622_6556<br>6715_7083 |
| IMR-L26 | 293-L12 | Novel | Late | 5622_6556 |
| IMR-L27 | 293-L13 | Novel | Late | 1585_1953 |
| IMR-L28 | NA | Novel | Late | 492_5530<br>5622_10636 |
| IMR-L29 | NA | Novel | Late | 492_5506<br>5622_10660 |
| IMR-L30 | NA | Novel | Late | 315_400 |
| IMR-L33 | 293-L21 | M5 | Late | 732_1426 |
| IMR-L60 | 293-L39 | M6 | Late | 732_1426<br>1585_1953 |
| NA | 293-L15 | Novel | Late | 492_5506<br>5622_6556<br>6715_10781 |

NA, not applicable.

### Wraparound transcripts are expressed from the late region of the JCPyV genome

Since wraparound transcripts, like the conserved late transcripts containing multiple copies of a leader sequence in diverse PyVs have been identified [25, 27–37], we aimed to elucidate their detailed structure in JCPyV. This study identified 15 late transcripts harboring a repeated sequence at their 5’ end (**Fig 4A and S3 Table**). These transcripts can be classified into two categories based on the repeating unit length: “377-WA” transcripts with a tandem repeat of 116 nt (nt position 377-492) up to three times and “401-WA” transcripts with a tandem repeat of 92 nt (nt position 401-492) up to four times (**Fig 4A**). Notably, in a subset of these wraparound transcripts, the region immediately upstream of the repeat (nt positions 299-376 in 377-WA and 299-400 in 401-WA, respectively) is spliced out (**Fig 4A**). The expression of these wraparound transcripts in IMR-32 and 293 cells was confirmed by reverse transcription (RT)-PCR using primers spanning the repetitive element, which yielded a ladder-like electrophoresis pattern (**Fig 4B and 4C**). Nanopore sequencing of these RT-PCR products confirmed the presence of PCR products responsible for leader-to-leader splicing (**S4 Table and S5 Table**). To evaluate the significance of these wraparound transcripts in pathology of JCPyV-related disease, we investigated the expression of the transcripts in brain tissues from PML patients where high copy number of JCPyV DNA was confirmed by real-time PCR (**S6 Table**). RT-PCR followed by nanopore sequencing analysis demonstrated that late wraparound transcripts were expressed in PML lesions although proportions of transcripts varied among cases (**Fig 4D, 4E, S7 Table, and S8 Table**). Collectively, these findings demonstrate the expression of wraparound transcripts from the late region of JCPyV genome in both cultured cells and PML lesions suggesting its contribution to pathology of PML.

**Fig 4:**
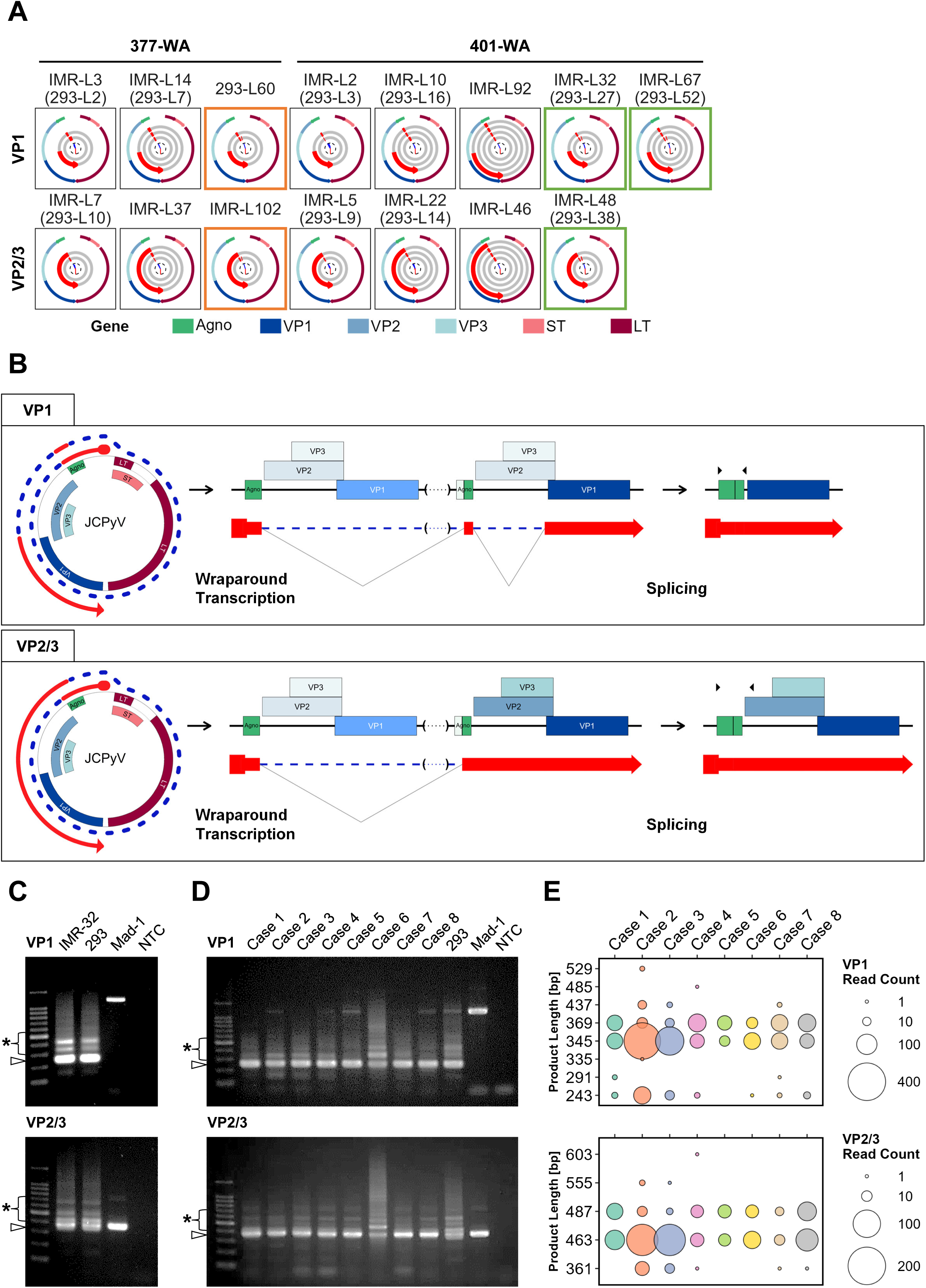
Wraparound transcripts encoded in the late region of the JCPyV genome. **A.** Watch plots illustrating wraparound transcripts from the late region. Histograms positioned at the center of the panels indicate the distribution of 5’ ends (blue) and 3’ ends (red) for each transcript. The exons of each transcript are represented as red segments, and arrows indicate the 5’ to 3’ direction. The transcripts are visualized from the innermost track to the outer track, progressing from the 5’ end to the 3’ end, starting from the most common 5’ ends and ending at the most common 3’ ends. For transcripts with overlapping exons, a panel contains multiple tracks. The outermost track displays viral ORFs. Transcripts are categorized based on the length of the repetitive element. Those with a repeating unit of 377-492 nt are designated as “377-WA,” and those with a repeating unit of 401-492 nt are labeled as “401-WA,” respectively. Transcripts lacking VP2/3 ORFs are shown in the upper row, while those with VP2/3 ORFs are depicted in the lower row. The region from nt position 299-376 is spliced out in transcripts outlined with an orange frame, and the region from nt position 299-400 is spliced out in transcripts outlined with a green frame. **B.** Schematic diagram illustrating wraparound transcripts from the late region. Arrowheads indicate the positions of primers used in RT-PCR (refer to **Fig 4C and 4D**). **C.** RT-PCR analysis detecting the late wraparound transcripts in cultured cells. The results of electrophoresis are shown. Arrowheads indicate PCR products representing non-wraparound transcripts, while asterisks indicate PCR products representing wraparound transcripts, respectively. **D.** RT-PCR analysis detecting the late wraparound transcripts in brain tissues from PML patients. The results of electrophoresis are shown. Arrowheads indicate PCR products representing non-wraparound transcripts, while asterisks indicate PCR products representing wraparound transcripts, respectively. **E.** Bubble charts visualizing read counts of the RT-PCR products from each PML case shown in **Fig 4D**. WA, wraparound; Mad-1, JCPyV Mad-1 strain genomic DNA; NTC, no template control.

### SuperT transcripts is conserved in JCPyV

The SuperT antigen is a splicing variant of the LT antigen, which contains a duplicated region with two copies of the LxCxE motif, a retinoblastoma (Rb) protein-binding motif. Since previous studies have demonstrated the conservation of SuperT transcripts in PyVs [25, 38], we explored SuperT transcripts in JCPyV. In this study, four early transcripts with a repetitive element containing multiple LxCxE motif (nt position 4,357-4,343) were identified by NGS (**Fig. 5A and S9 Table**), although the abundance of sequencing reads corresponding to these transcripts was very low (**S9 Table**). These transcripts are characterized by a tandem repeat of 153 nt (nt position 4,425-4,273) with repeat number ranged from three to nine (**Fig. 5A**). Expression of these wraparound transcripts in IMR-32 and 293 cells were validated by RT-PCR with primers which amplify the repetitive element, resulting in a ladder-like electrophoresis pattern (**Fig. 5B**, **Fig. 5C**). DNA sequences of these RT-PCR products were analyzed by nanopore sequencing, detecting wraparound-specific sequences (**S10 Table**). Nanopore sequencing following RT-PCR analysis also demonstrated expression of SuperT transcripts in brain tissues from PML patients with an inverse correlation between transcript length and expression level (**Fig 5D, 5E and S11 Table**). Taken together, these findings demonstrate the expression of SuperT transcripts from the early region of JCPyV genome in both cultured cells and PML lesions suggesting its contribution to pathology of PML.

**Fig 5:**
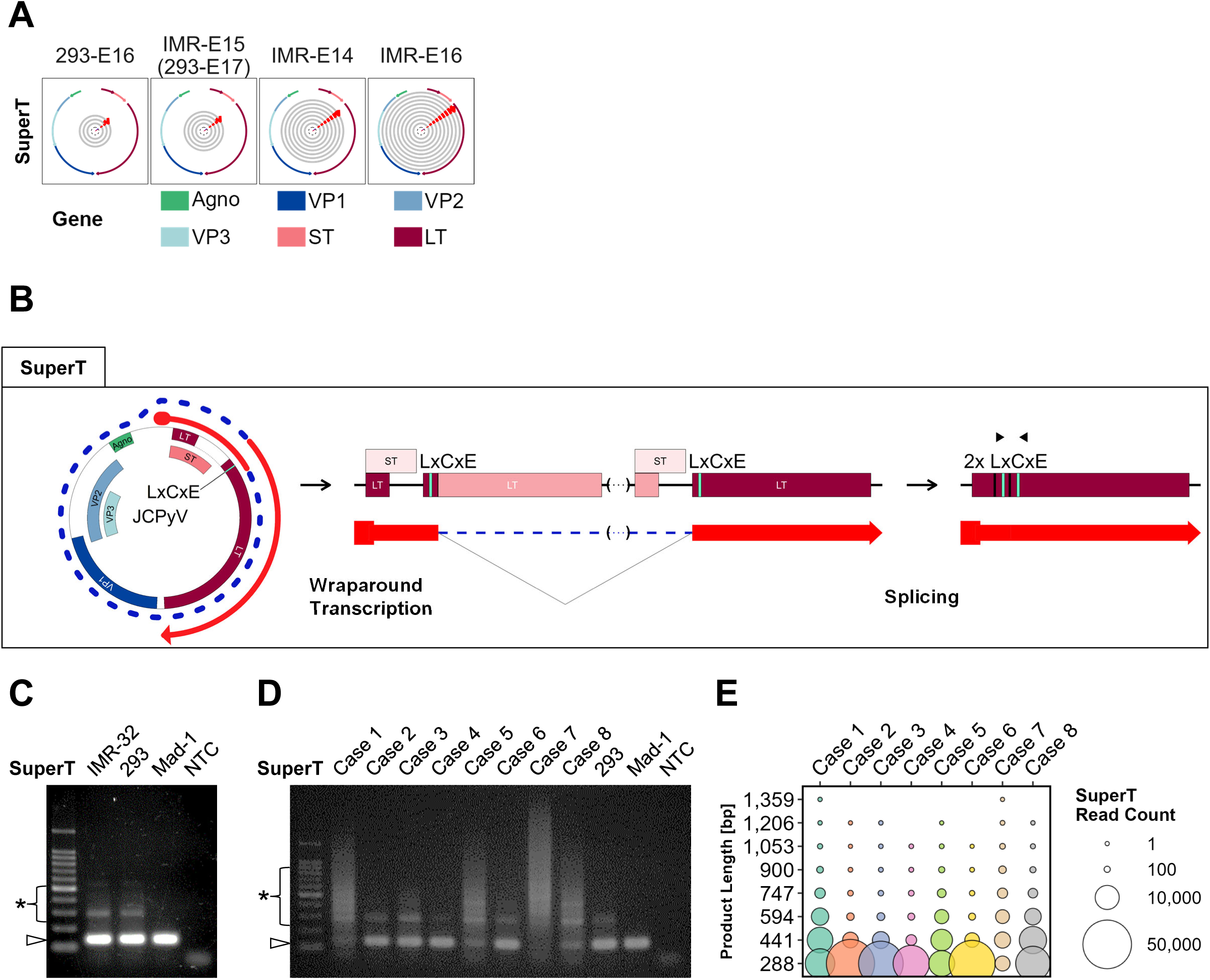
SuperT transcripts are conserved in JCPyV. **A.** Watch plots illustrating SuperT transcripts in JCPyV, presented in a similar manner to that described in Fig 4. **B.** Schematic diagram displaying the SuperT transcript from the early region. Arrowheads indicate the positions of primers used in RT-PCR (refer to **Fig 5C and 5D**) and green lines indicate LxCxE motifs. **C.** RT-PCR analysis detecting SuperT transcripts in cultured cells. The result of electrophoresis is shown. Arrowheads indicate PCR products representing non-wraparound transcripts, while asterisks indicate PCR products representing SuperT transcripts, respectively. **D.** RT-PCR analysis detecting SuperT transcripts in brain tissues from PML patients. The result of electrophoresis is shown. Arrowheads indicate PCR products representing non-wraparound transcripts, while asterisks indicate PCR products representing SuperT transcripts, respectively. **E.** Bubble charts visualizing read counts of the RT-PCR products from each PML case shown in **Fig 5D**. Mad-1, JCPyV Mad-1 strain genomic DNA; NTC, no template control.

### Expression pattern of wraparound transcripts is not affected by alteration in the NCCR sequence

Since rearrangements in the NCCR can alter viral gene expression [39, 40], we hypothesized that NCCR rearrangements could affect the expression pattern of wraparound transcripts. To examine this hypothesis, we determined the whole genome sequence of JCPyV in PML cases by amplicon sequencing and analyzed the NCCR structure. Our analysis revealed that JCPyV in all PML cases possessed the PML-type NCCR with complex rearrangements as well as nucleotide substitutions, insertions, and deletions (**Fig 6 and S7 Fig**). Although the proportion of the late wraparound (VP1 and VP2/3) transcripts in cases 2 and 3 was distinct from that in other cases (**Fig 4E**), and the proportion of early (SuperT) transcripts in case 7 was distinct from that in other cases (**Fig 5E**), significant variation in the NCCR structure was not observed in these cases (**Fig 6 and S7 Fig**). These data suggest that the NCCR does not significantly regulate the expression of wraparound transcripts in JCPyV.

**Fig 6:**
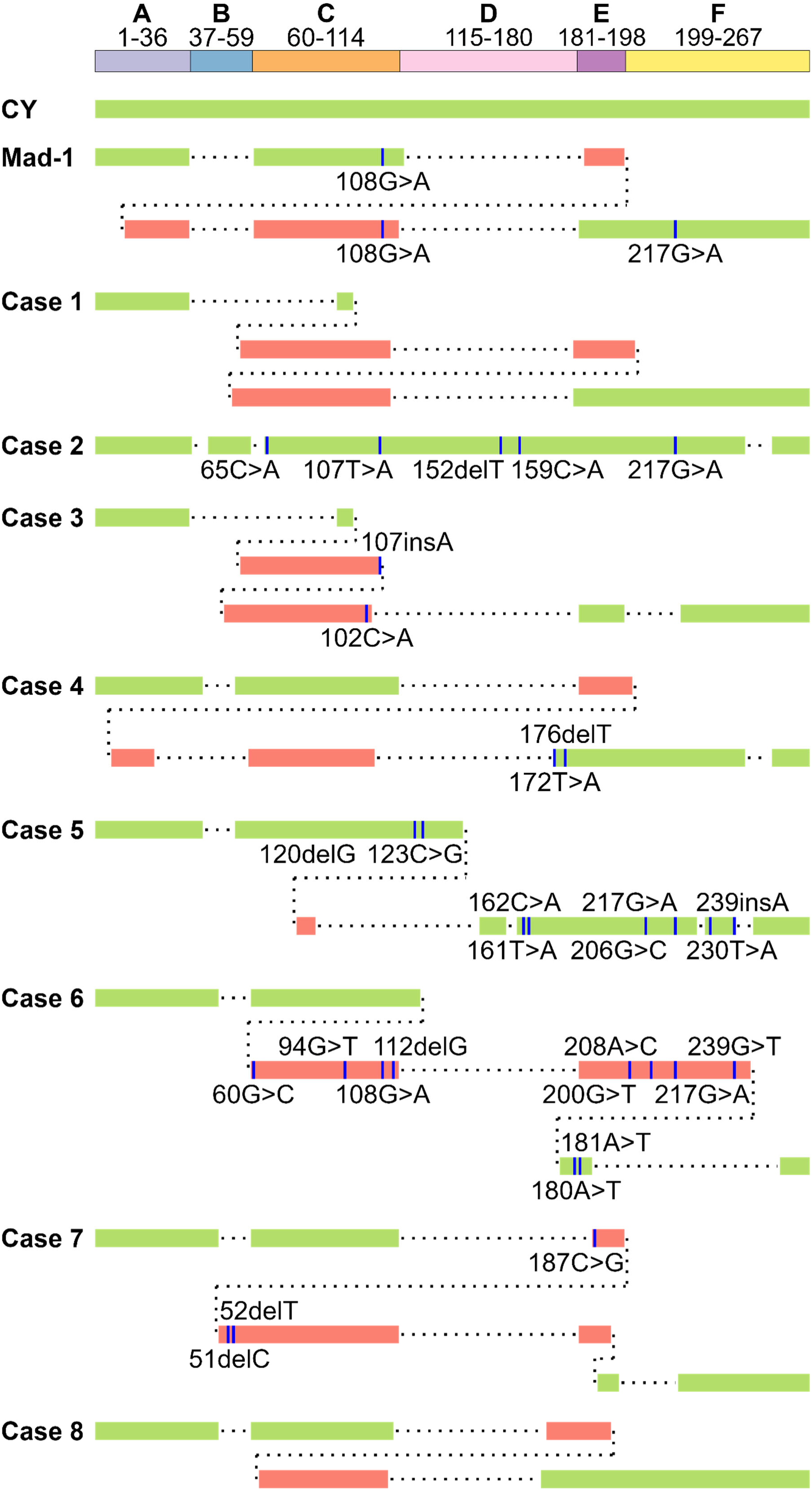
Structure of the NCCR of the JCPyV genome in PML cases. Structure of the NCCR, determined by amplicon sequencing, is shown. Both green and red lines represent components of the NCCR, while dashed lines indicate missing regions. Nucleotide substitutions, insertions, and deletions are shown as blue lines. Each number indicates the position in the JCPyV CY strain genome. The NCCRs of the archetype (CY) and PML-type (Mad-1) JCPyVs are also shown.

## Discussion

Long-read sequencing techniques, such as SMRTseq and nanopore sequencing, have revolutionized our understanding of viral transcriptomes, revealing complexities previously underestimated [24]. A recent study combining long-read and short-read sequencing elucidated the detailed viral transcriptome of SV40 and BKPyV [25]. While previous reports have characterized JCPyV transcripts using non-NGS methods, a comprehensive understanding was lacking. In this study, we combined long-read and short-read sequencing to comprehensively analyze the JCPyV transcriptome, identifying 39 novel transcripts, including wraparound transcripts (**Fig 2 and Fig 3**). To our knowledge, this represents the first study to provide a comprehensive catalog of JCPyV transcripts using long-read sequencing technology.

We identified 15 wraparound transcripts transcribed from the late region of the JCPyV genome, characterized by a leader-to-leader sequence (**Fig 4**). Similar transcripts, initially discovered in murine PyV [27], have been shown to play a critical role in both late gene mRNA expression and viral replication [31, 33]. A recent study confirmed the conservation of the leader-to-leader junction of late transcripts among various PyVs [25]. These findings suggest that late wraparound transcripts may be essential for PyV infection. However, their precise role in the gene expression and replication of JCPyV remains unclear, necessitating further functional analyses. The LT antigen is pivotal in PyV replication, primarily through its interaction with the Rb protein via the LxCxE motif, leading to cell cycle dysregulation [41, 42]. SuperT, a splicing variant of the LT antigen with tandem repeats, was originally discovered in SV40-transformed cells [38]. While a previous study demonstrated conservation of the SuperT transcript among PyVs and identified a SuperT-specific splicing junction in JCPyV using short-read sequencing [25], the full-length sequence remained unknown. Our long-read sequencing analysis revealed that JCPyV expresses SuperT transcripts with multiple LxCxE motifs (**Fig 5**). The increased number of these motifs might enhance the Rb binding capacity of SuperT, making it a more potent Rb suppressor compared to the LT antigen. The role of SuperT in JCPyV biology and the pathogenesis of JCPyV-related diseases requires further analysis.

JCPyV exhibits a complex array of mRNAs utilizing diverse splicing patterns (**Fig 3**). In this study, we identified several novel splice sites, including a site at position 400 of the JCPyV genome that serves as both a splicing acceptor and donor. Dual utilization of splice sites, broadening the diversity of splicing variants, was first identified in the human interferon regulatory factor-3 (IRF-3) gene [43]. A subsequent study have found numerous dual-function splice sites across human and mouse genomes [44]. In our study, the dual-specific splice site contributes to generating 13 distinct mRNA species among the 39 novel transcripts identified (**Fig 3**), highlighting its role in expanding the transcriptomic complexity of the compact JCPyV genome.

PML is characterized by demyelination caused by lytic infection of JCPyV to oligodendrocytes under immunosuppressive conditions [45]. JCPyV isolated from PML lesions often exhibits genomic alterations [46–51]. In particular, rearrangements of the NCCR, transitioning from archetype to PML-type, play a critical role in PML development by altering viral gene expression, replication efficiency, and cellular tropism [39, 40, 45]. In addition to viral genomic analysis, a previous study reported upregulation of human genes associated with neuroinflammation, blood-brain barrier permeability, and neurodegenerative diseases was found in PML brains, as determined by short-read sequencing techniques [52]. Although these studies focused on genomic alterations of the virus or host gene expression, little is known about viral gene expression in PML. Our data demonstrates the expression of novel viral wraparound transcripts in PML tissues, with variation in wraparound transcript proportions among cases, suggesting differential viral mRNA expression through PML development (**Fig 4 and Fig 5**). The significance of these transcripts in PML pathogenesis and their regulatory mechanisms warrant further investigation.

This study has several limitations. Firstly, we used only the Mad-1 strain of JCPyV to investigate viral transcripts, and the comprehensive viral transcriptome was not determined in PML cases. Consequently, some of the novel transcripts identified may be specific to the Mad-1 strain. Secondly, our experimental approach utilized cultured cells transfected with JCPyV DNA instead of naturally infected cells, potentially affecting the expression pattern of viral transcripts. Thirdly, the sample size of PML was limited to only eight cases. This limited number of clinical specimens may introduce sampling bias, potentially affecting the generalizability of our findings. Further analysis involving a larger cohort of PML cases is necessary to validate and extend the results of the present study. Lastly, despite performing whole-genome sequencing of JCPyV in PML cases, we did not find a clear correlation between the wraparound transcript expression patterns and PML-type NCCR structure.

In conclusion, our study identified several novel JCPyV transcripts through combined short-read and long-read sequencing approaches, significantly enhancing our understanding of the JCPyV transcriptome. Additionally, we demonstrated the expression of SuperT and wraparound transcripts in PML tissues. While further studies are necessary to elucidate their biological functions and protein-coding potential, particularly in the context of PML pathogenesis, our findings provide valuable insights into the molecular basis of JCPyV biology and PML development. This work underscores the potential of long-read sequencing for investigating viral transcriptomics and may contribute to developing novel countermeasures against PML in the future.

## Materials and Methods

### Cell culture

Human neuroblastoma cells (IMR-32) were cultured in Dulbecco’s Modified Eagle Medium (DMEM, Thermo Fisher Scientific, Waltham, MA) supplemented with 10% fetal bovine serum (FBS) and MEM non-essential amino acids solution (Thermo Fisher Scientific). Human embryonic kidney cells (293) were grown in DMEM supplemented with 5% FBS. Both cell lines were incubated at 37°C with 5% CO_2_.

### Preparation of circular viral genomic DNA

The complete genome of JCPyV Mad-1 strain (NCBI RefSeq accession number NC_001699.1) containing an insertion of GGTC between nt positions 109 and 110 was subcloned into the BamHI site of the pUC19 vector [50]. The plasmid was digested with BamHI and self-ligated to generate a complete circular JCPyV genome.

### Viral genome transfection and RNA extraction

IMR-32 cells were transfected with circular JCPyV genomic DNA as described previously [50]. Briefly, 1×10^5^ cells were seeded onto type I collagen-coated 24-well plates and transfected with 200 ng of viral genome using Attractene Transfection Reagent (Qiagen, Hilden, Germany). One day post-transfection, the cells were transferred to type I collagen-coated 6-well plates and cultured for an additional three days before harvesting. For 293 cells, 6×10^5^ cells were seeded onto type I collagen-coated 6-well plates and transfected with 200 ng of viral DNA using Attractene Transfection Reagent (Qiagen). Two days later, the cells were transferred to type I collagen-coated 60 mm dishes and maintained for an additional day before collection. Total RNA was extracted from all cells using Isogen reagent (Nippon Gene, Tokyo, Japan).

### Quality check of RNA

Total RNA concentration was quantified using Qubit 4 Fluorometer (Thermo Fisher Scientific) with the Qubit RNA High Sensitivity Assay Kit (Thermo Fisher Scientific). RNA quality was assessed using 2100 Bioanalyzer (Agilent, Santa Clara, CA) with the RNA 6000 Pico Kit (Agilent). RNA samples with RNA integrity number values greater than 9.0 were subjected to downstream analysis.

### Short-read RNA sequencing

Total RNA was either enriched for poly(A)+ RNA or depleted of ribosomal RNA (rRNA) prior to construction of short-RNAseq libraries. Poly(A)+ RNA was isolated from 1 μg of total RNA using the NEBNext Poly(A) mRNA Magnetic Isolation Module (New England Biolabs, Ipswich, MA), whereas rRNA-depleted RNA was prepared from 1 μg of total RNA using the NEBNext rRNA Depletion Kit v2 (Human/Mouse/Rat) (New England Biolabs). NGS libraries were generated from poly(A)+ RNA or rRNA-depleted RNA using the NEBNext Ultra II Directional RNA Library Prep Kit for Illumina (New England Biolabs) and NEBNext Multiplex Oligos for Illumina (96 Unique Dual Index Primer Pairs) (New England Biolabs), following the manufacturer’s instructions. Each library was sequenced on NextSeq 1000 platform (Illumina, San Diego, CA) using the NextSeq 1000/2000 P2 Reagents (300 Cycles) v3 (Illumina) to generate 150 base pairs paired-end reads. Prior to read mapping, FASTQ files were processed using the Nextflow software (version 23.04.1) [53] with process_illumina.nf pipeline, as previously described [25]. Briefly, both read 1 (R1) and read 2 (R2) were trimmed with Trim Galore software (version 0.6.10). R1 sequences were then reverse-complemented, and all reads in both R1 and R2 files were labelled with “_1” or “_2” suffixes, respectively. Finally, R1 and R2 files were concatenated. The processed FASTQ files were then subjected to read mapping described below.

### Direct RNA sequencing

Total RNA (1 μg) was used to purify poly(A)+ RNA according to the method described above. Sequencing libraries were generated from 50 ng of poly(A)+ RNA using the Direct RNA Sequencing Kit (catalog number SQK-RNA002, Oxford Nanopore Technologies, Oxford, UK). Sequencing was performed on a MinION device using an R9.4.1 flow cell (Oxford Nanopore Technologies) for up to 20 hours. Guppy software (version 4.4.1) was used for base calling, converting FAST5 files to FASTQ files. The following command was employed: guppy_basecaller -i fast5 -s fastq --flowcell FLO-MIN106 --kit SQK-RNA002 -r --trim_strategy rna --reverse_sequence true --u_substitution true. Subsequently, the FASTQ files were subjected to the read mapping procedure described below.

### Single-molecule real-time sequencing

SMRTseq was conducted by Macrogen, Inc (Seoul, Republic of Korea). IsoSeq libraries were constructed from poly(A)+ RNA using the SMRTbell prep kit 3.0 (Pacific Biosciences, Menlo Park, CA). Each sample was sequenced individually on SMRT Cell 8M (Pacific Biosciences) with Sequel II system (Pacific Biosciences). HiFi reads were refined using Isoseq3 software (version 3.8.2) and subsequently converted to FASTQ files using BAM2fastx software (version 3.0.0). The generated FASTQ files were then subjected to the read mapping process described below.

### Read mapping

NGS data analysis was performed using the main.nf Nextflow pipeline, following the methodology described in a previous study [25]. Sequencing reads generated by short-RNAseq (pre-processed with the process_illumina.nf pipeline), dRNAseq, and SMRTseq were aligned to JCPyV genome (NCBI RefSeq accession number NC_001699.1) using minimap2 software (version 2.26) [54, 55], generating BAM files. For short-RNAseq, the reference sequence consisted of two concatenated copies of JCPyV genome. In contrast, sequencing reads from dRNAseq and SMRTseq were mapped to 20 copies of concatenated JCPyV genome. Introns were identified by minimap2 and labeled with the “N” CIGAR flag. Subsequently, BAM files were converted to BED files using bedtools software (version 2.31.0) [56]. Importantly, due to the presence of multiple genome copies in the reference sequence, reads were randomly assigned to the viral genomes in the BED files. To address this, all reads were aligned to the genome copy located at the 5’ end of the reference sequence using bed_slide_wraparound_reads.py. Finally, the BED files were processed with bed_to_span.py to generate spans files.

### Calculation of sequencing depth

To calculate sequencing depth, the BAM files generated by minimap2 were processed using the main.nf Nextflow pipeline described above. This pipeline employed a similar approach as used for BED files, aligning the sequencing reads in the BAM files to the genomic copy located at the 5’ end of the reference sequence using bam_slide_wraparound_reads.py. With the aid of samtools software (version 1.17) [57], the sequencing reads were segregated into forward and reverse strands, enabling the calculation of sequencing depth for each strand. The read depth was expressed as a relative value, with the highest value on the positive/negative strand normalized to 1.

### Identification of viral transcripts

During spans file generation with bed_to_span.py, transcript classes were defined based on shared intron combinations. This means sequencing reads with identical sets of introns were grouped into the same group. Consequently, start and end positions of a transcript can vary within a transcript class. For dRNAseq and SMRTseq data, only transcript classes where all introns were supported by at least five short-RNAseq (rRNA-) reads spanning the splicing junction were retained for further analysis. Within each strand, transcript class numbers were assigned based on the relative abundance of reads, with lower numbers indicating higher read counts. Transcript IDs were constructed by concatenating the cell of detection (IMR-32 or 293), the kinetic class (early or late), and the assigned transcript class number. In this study, only transcripts exceeding 0.1% of reads from each strand in dRNAseq or SMRTseq were visualized. However, exceptions were made for previously identified transcripts, including M5 and M6 late transcripts, and SuperT which were also validated by RT-PCR. The names of known transcripts were adopted from previous studies [18, 21–23].

### Reverse transcription-PCR

Repeated sequences in wraparound transcripts were amplified from 50 pg of poly(A)+ RNA, purified as described above, by RT-PCR using QIAGEN OneStep RT-PCR Kit (Qiagen) with following primers (5’ to 3’): VP1 wraparound transcripts forward primer (JCV_269_F) AGCTGGCCATGGTTCTTCG and reverse primer (JCV_1455_R) AAGAGCAGGTGTTACAGTCC, VP2/3 wraparound transcripts forward primer (JCV_269_F) AGCTGGCCATGGTTCTTCG and reverse primer (JCV_639_R) TATAGTAGCAGCAGCCTCTC, and SuperT transcript forward primer (JCV_4287_F) TTAGGTGGGGTAGAGTGTTG and reverse primer (JCV_4421_R) AACCTATGGAACAGATGAATGG. PCR products were separated on 2% agarose gel and stained with ethidium bromide.

### Amplicon sequencing

Nanopore amplicon sequencing was employed to detect DNA sequences corresponding with wraparound transcripts from each RT-PCR product (refer to the “Reverse transcription-PCR” section described above). PCR products were purified using GEL/PCR Purification Column (Favorgen, Ping Tung, Taiwan). The purified products then underwent library construction with Ligation Sequencing Kit (catalog number SQK-LSK109, Oxford Nanopore Technologies) and Native Barcoding Expansion (catalog number EXP-NBD104 and EXP-NBD114, Oxford Nanopore Technologies) according to the manufacturer’s instructions. Sequencing was performed on an R9.4.1 flow cell using a MinION device (Oxford Nanopore Technologies). Base calling was conducted with Guppy software (version 4.4.1) using the following command: guppy_basecaller -i fast5 -s fastq --flowcell FLO-MIN106 --kit SQK-LSK109 -- barcode_kits EXP-NBD104 (or EXP-NBD114) --trim_barcodes –q 100000. Read mapping was carried out using methods similar to those described for dRNAseq above. Total read counts were calculated using samtools software (version 1.17).

### Data visualization

NGS data, converted to spans format, was visualized using R software (version 4.3.2) with modifications to the codes published previously [25].

### Data and code availability

All raw sequencing data are registered in FASTQ format under BioProject accession number PRJDB18771 in the Sequence Read Archive database of the DNA Data Bank of Japan. Nextflow pipelines, along with the associated bash and python scripts used for NGS data processing, and R scripts for data visualization were developed by Nomburg et al (https://github.com/jnoms/SV40_transcriptome). R codes used for overview of the JCPyV transcripts (**Fig 3A**), bubble chart (**Fig 4E and 5E**), and visualization of NCCR (**Fig 6**) are available on GitHub (https://github.com/iidashun/JCPyV_transcriptome).

### Clinical samples

All procedures involving clinical samples were carried out with the approval of the Institutional Review Board of the National Institute of Infectious Diseases (approval number 1746). In this study, eight individuals diagnosed histopathologically with PML were enrolled. Frozen brain tissues from these patients were analyzed for the presence of JCPyV wraparound transcripts and viral DNA. In brief, total RNA was extracted from the brain tissues using Isogen reagent (Nippon Gene) and treated with DNase I (Zymo Research, Irvine, CA). Total RNA was then subjected to RT-PCR and amplicon sequencing to detect repeating units of wraparound transcripts as described above. For viral DNA detection, DNA was purified from brain tissues using DNeasy Blood & Tissue Kit (Qiagen). Viral DNA was quantified by TaqMan real-time PCR [58] using Thunderbird Probe qPCR Mix (Toyobo, Osaka, Japan). Human β-Actin DNA was also quantified by TaqMan real-time PCR as a endogenous control [59].

### Virus genome sequencing

The whole genome sequence of JCPyV was determined by amplicon sequencing using a Nanopore sequencer with library construction using the Ligation Sequencing Kit (catalog number SQK-LSK109, Oxford Nanopore Technologies). Briefly, DNA was extracted from PML brain tissues using DNeasy Blood & Tissue Kit (Qiagen). The JCPyV genome was amplified by multiplex PCR using Q5 Hot Start High-Fidelity 2X Master Mix (New England Biolabs) and primers in two different pools (**S12 Table**). PCR products from pools 1 and 2 were mixed, cleaned up, and then treated with NEBNext Ultra II End Repair/dA-Tailing Module (New England Biolabs). The PCR products were subjected to barcode ligation using Native Barcoding Expansion (catalog number EXP-NBD104 and EXP-NBD114, Oxford Nanopore Technologies) and Blunt/TA Ligase Master Mix (New England Biolabs), followed by adapter ligation using NEBNext Quick Ligation Module (New England Biolabs). Sequencing was performed on a MinION device using an R9.4.1 flow cell (Oxford Nanopore Technologies). Guppy (version 4.4.1) and Porechop (version 0.2.4) [60] software were used for base calling and adapter/primer trimming, respectively. Sequencing reads were mapped to the JCPyV genome (NCBI RefSeq accession number NC_001699.1) using CLC Genomics Workbench (Qiagen), and the resultant consensus sequences were analyzed manually to detect alterations of DNA sequences in the NCCR region, comparing them with the NCCR region of archetype JCPyV (CY strain, NCBI GenBank accession number AB038249.1).

## Acknowledgements

We specially thank Dr. Yasuko Orba and Dr. Hirofumi Sawa of Hokkaido University for their kind gift of the JCPyV genome-coding vector.

## Supporting Information

**S1 Table:** Read counts from each sequencing method in IMR-32 cells.

| Method | Total reads (%) | Mapped reads (%) | Spliced reads (%) |
| --- | --- | --- | --- |
| Short-RNAseq [poly(A)+] | 100,412,315 | 3,558,198 | 140,719 |
|  | (100.00) | (3.54) | (0.14) |
| Short-RNAseq [rRNA-] | 103,727,302 | 2,629,019 | 74,016 |
|  | (100.00) | (2.53) | (0.07) |
| dRNAseq | 555,695 | 3,284 | 1,028 |
|  | (100.00) | (0.59) | (0.18) |
| SMRTseq | 3,430,793 | 67,848 | 31,237 |
|  | (100.00) | (1.98) | (0.91) |

Short-RNAseq, short-read RNA sequencing; rRNA, ribosomal RNA; dRNAseq direct RNA sequencing; SMRTseq, single-molecule real-time sequencing.

**S2 Table:** Read counts from each sequencing method in 293 cells.

| Method | Total reads (%) | Mapped reads (%) | Spliced reads (%) |
| --- | --- | --- | --- |
| Short-RNAseq [poly(A)+] | 111,323,204 | 2,106,501 | 131,863 |
|  | (100.00) | (1.89) | (0.12) |
| Short-RNAseq [rRNA-] | 113,886,191 | 1,798,614 | 67,947 |
|  | (100.00) | (1.58) | (0.06) |
| dRNAseq | 228,666 | 389 | 218 |
|  | (100.00) | (0.17) | (0.10) |
| SMRTseq | 2,317,212 | 34,028 | 24,156 |
|  | (100.00) | (1.47) | (1.04) |

Short-RNAseq, short-read RNA sequencing; rRNA, ribosomal RNA; dRNAseq direct RNA sequencing; SMRTseq, single-molecule real-time sequencing.

**S3 Table:** Wraparound transcripts encoded in the late region of the JCPyV genome.

| ID (IMR-32) | ID (293) | Early/Late | ORF | Introns | Percentage<br>in IMR-32,<br>dRNAseq<br>(%) | Percentage<br>in IMR-32,<br>SMRTseq<br>(%) | Percentage<br>in 293,<br>dRNAseq<br>(%) | Percentage<br>in 293,<br>SMRTseq<br>(%) |
| --- | --- | --- | --- | --- | --- | --- | --- | --- |
| IMR-L2 | 293-L3 | Late | VP1 | 492_5530<br>5622_6556 | 13.37 | 0.20 | 7.32 | 0.05 |
| IMR-L3 | 293-L2 | Late | VP1 | 492_5506<br>5622_6556 | 8.36 | 0.20 | 8.78 | 0.10 |
| IMR-L10 | 293-L16 | Late | VP1 | 492_5530<br>5622_10660<br>10752_11686 | 0.98 | 0.03 | 0.49 | UDL |
| IMR-L14 | 293-L7 | Late | VP1 | 492_5506<br>5622_10636<br>10752_11686 | 0.49 | 0.01 | 1.46 | 0.01 |
| IMR-L32 | 293-L27 | Late | VP1 | 298_400<br>492_5530<br>5622_6556 | 0.10 | 0.05 | UDL | 0.03 |
| IMR-L67 | 293-L52 | Late | VP1 | 298_400<br>492_5530<br>5622_10660<br>10752_11686 | UDL | 0.01 | UDL | 0.00 |
| IMR-L92 | NA | Late | VP1 | 492_5530<br>5622_10660<br>10752_15790<br>15882_16816 | UDL | 0.00 | UDL | UDL |
| NA | 293-L60 | Late | VP1 | 298_376<br>492_5506<br>5622_6556 | UDL | UDL | UDL | 0.00 |
| IMR-L5 | 293-L9 | Late | VP2/3 | 492_5530 | 4.42 | 0.26 | 0.98 | 0.05 |
| IMR-L7 | 293-L10 | Late | VP2/3 | 492_5506 | 1.87 | 0.16 | 0.98 | 0.04 |
| IMR-L22 | 293-L14 | Late | VP2/3 | 492_5530<br>5622_10660 | 0.29 | 0.03 | 0.49 | UDL |
| IMR-L37 | NA | Late | VP2/3 | 492_5506<br>5622_10636 | 0.10 | 0.01 | UDL | UDL |
| IMR-L46 | NA | Late | VP2/3 | 492_5530<br>5622_10660<br>10752_15790 | 0.10 | UDL | UDL | UDL |
| IMR-L48 | 293-L38 | Late | VP2/3 | 298_400<br>492_5530 | UDL | 0.05 | UDL | 0.01 |
| IMR-L102 | NA | Late | VP2/3 | 298_376<br>492_5506 | UDL | 0.00 | UDL | UDL |

ORF, open reading frame; NA, not applicable; UDL, under detection limit.

**S4 Table:** Read counts from amplicon sequencing detecting JCPyV late (VP1) wraparound ripts.

| Product length (bp) | Introns | ID (IMR-32) | ID (293) | Read count (IMR-32) | Read count (293) |
| --- | --- | --- | --- | --- | --- |
| 529 | 492_5530 | IMR-L92 | NA | 1 | 0 |
|  | 5622_10660 |  |  |  |  |
|  | 10752_15790 |  |  |  |  |
|  | 15882_16816 |  |  |  |  |
| 485 | 492_5506 | IMR-L14 | 293-L7 | 2 | 12 |
|  | 5622_10636 |  |  |  |  |
|  | 10752_11686 |  |  |  |  |
| 437 | 492_5530 | IMR-L10 | 293-L16 | 36 | 6 |
|  | 5622_10660 |  |  |  |  |
|  | 10752_11686 |  |  |  |  |
| 369 | 492_5506 | IMR-L3 | 293-L2 | 261 | 389 |
|  | 5622_6556 |  |  |  |  |
| 345 | 492_5530 | IMR-L2 | 293-L3 | 442 | 214 |
|  | 5622_6556 |  |  |  |  |
| 335 | 298_400 | IMR-L67 | 293-L52 | 4 | 1 |
|  | 492_5530 |  |  |  |  |
|  | 5622_10660 |  |  |  |  |
|  | 10752_11686 |  |  |  |  |
| 243 | 298_400 | IMR-L32 | 293-L27 | 41 | 16 |
|  | 492_5530 |  |  |  |  |
|  | 5622_6556 |  |  |  |  |

bp, base pairs; NA, not applicable.

**S5 Table:**
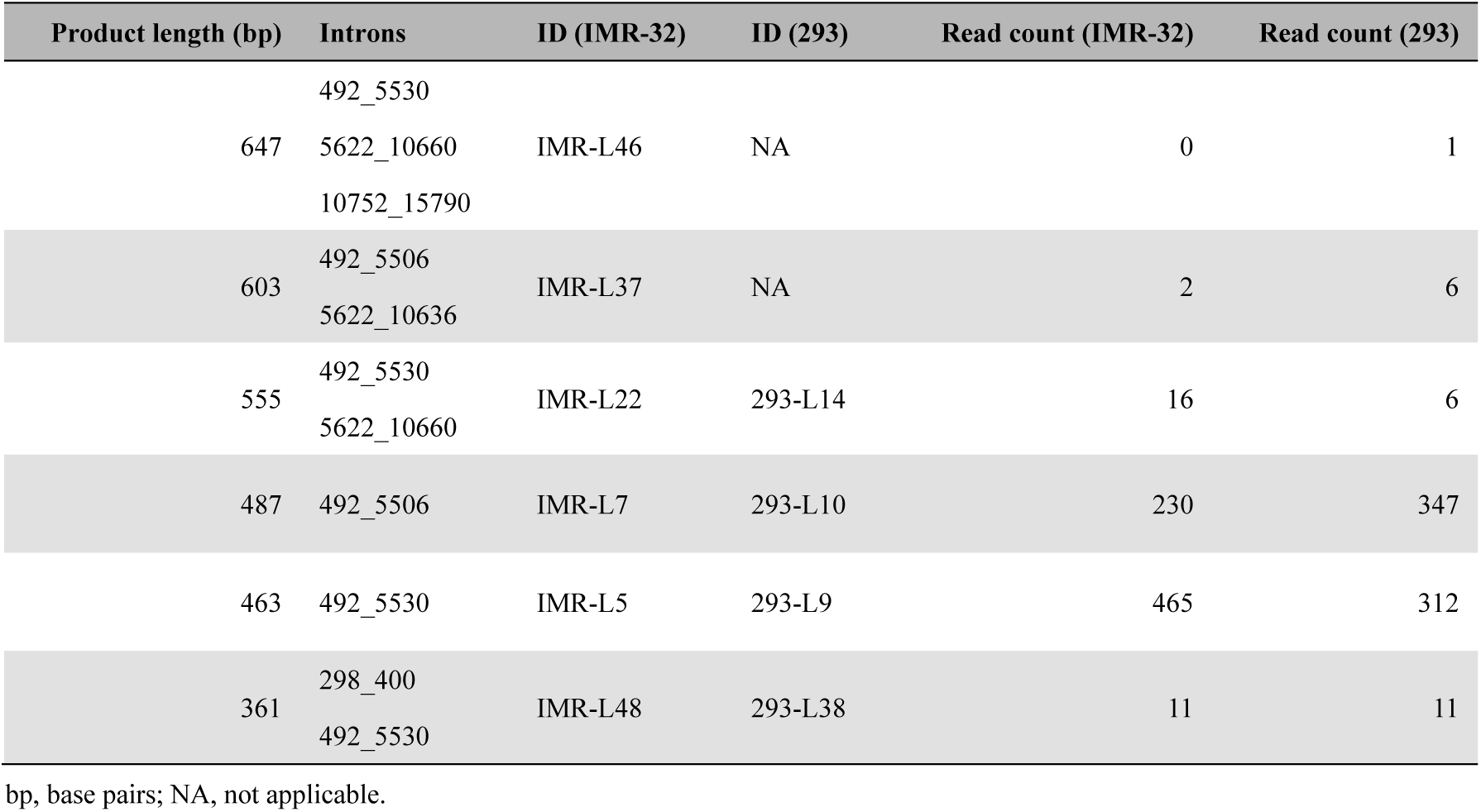
Read counts from amplicon sequencing detecting JCPyV late (VP2/3) wraparound ripts.

**S6 Table:** The list of PML cases.

| Case | JCPyV (copies/ng DNA) | β-actin (copies/ng DNA) | JCPyV (copies/cell) |
| --- | --- | --- | --- |
| 1 | $1.59 \times 10^6$ | $4.82 \times 10^2$ | $6.62 \times 10^3$ |
| 2 | $1.94 \times 10^6$ | $5.66 \times 10^2$ | $6.83 \times 10^3$ |
| 3 | $2.36 \times 10^6$ | $4.60 \times 10^2$ | $1.02 \times 10^4$ |
| 4 | $1.88 \times 10^6$ | $5.70 \times 10^2$ | $6.58 \times 10^3$ |
| 5 | $1.23 \times 10^6$ | $3.74 \times 10^2$ | $6.58 \times 10^3$ |
| 6 | $6.26 \times 10^6$ | $3.37 \times 10^2$ | $3.72 \times 10^4$ |
| 7 | $1.36 \times 10^6$ | $6.97 \times 10^2$ | $3.91 \times 10^3$ |
| 8 | $1.76 \times 10^5$ | $1.61 \times 10^1$ | $2.20 \times 10^4$ |

**S7 Table:**
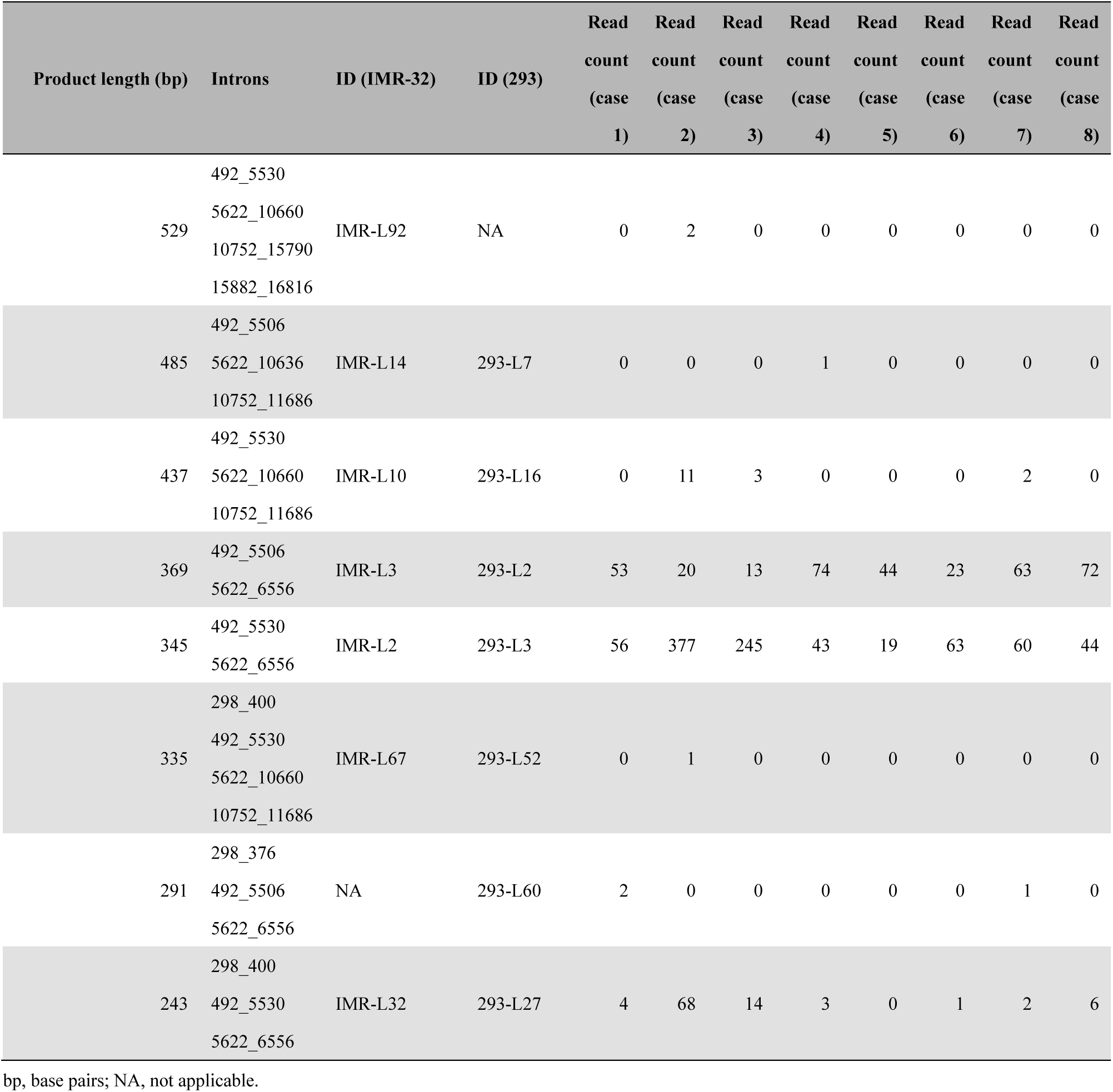
Read counts from amplicon sequencing detecting JCPyV late (VP1) wraparound ripts from PML samples.

**S8 Table:** Read counts from amplicon sequencing detecting JCPyV late (VP2/3) wraparound ripts from PML samples.

| Product length (bp) | Introns | ID (IMR-32) | ID (293) | Read count (case 1) | Read count (case 2) | Read count (case 3) | Read count (case 4) | Read count (case 5) | Read count (case 6) | Read count (case 7) | Read count (case 8) |
| --- | --- | --- | --- | --- | --- | --- | --- | --- | --- | --- | --- |
| 603 | 492_5506<br>5622_10636 | IMR-L37 | NA | 0 | 0 | 0 | 1 | 0 | 0 | 0 | 0 |
| 555 | 492_5530<br>5622_10660 | IMR-L22 | 293-L14 | 0 | 2 | 1 | 0 | 0 | 0 | 0 | 0 |
| 487 | 492_5506 | IMR-L7 | 293-L10 | 30 | 9 | 3 | 16 | 18 | 17 | 6 | 42 |
| 463 | 492_5530 | IMR-L5 | 293-L9 | 46 | 135 | 135 | 19 | 15 | 35 | 10 | 62 |
| 361 | 298_400<br>492_5530 | IMR-L48 | 293-L38 | 0 | 21 | 8 | 1 | 0 | 0 | 1 | 1 |

bp, base pairs; NA, not applicable.

**S9 Table:**
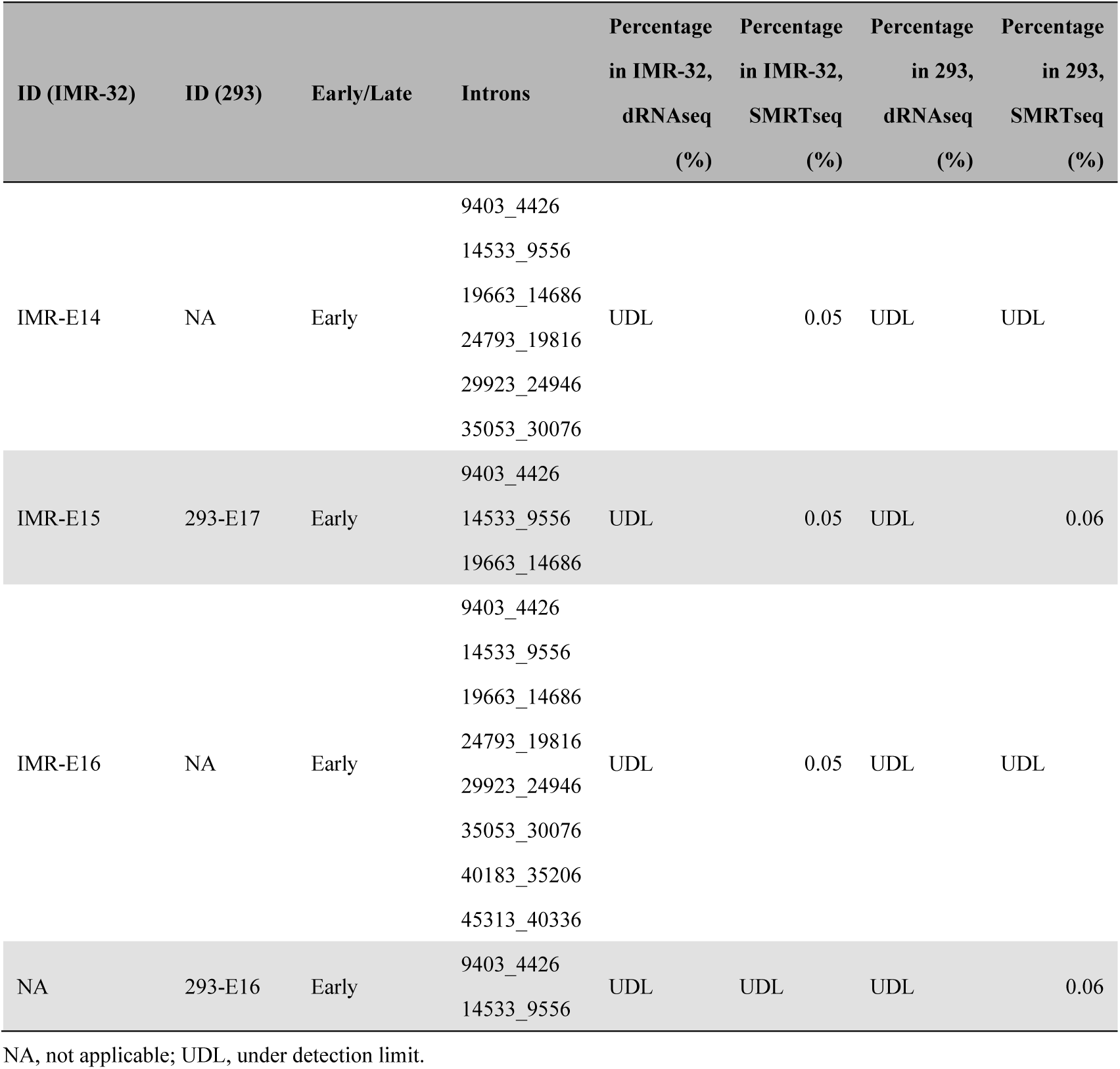
Wraparound transcripts encoded in the early region of the JCPyV genome.

**S10 Table:** Read counts from amplicon sequencing detecting JCPyV early wraparound transcripts.

| Product length (bp) | Introns | ID (IMR-32) | ID (293) | Read count (IMR-32) | Read count (293) |
| --- | --- | --- | --- | --- | --- |
| 747 | 9403_4426 | NA | NA | 11 | 3 |
|  | 14533_9556 |  |  |  |  |
|  | 19663_14686 |  |  |  |  |
|  | 24793_19816 |  |  |  |  |
| 594 | 9403_4426 | IMR-E15 | 293-E17 | 102 | 146 |
|  | 14533_9556 |  |  |  |  |
|  | 19663_14686 |  |  |  |  |
| 441 | 9403_4426 | NA | 293-E16 | 2,010 | 2,058 |
|  | 14533_9556 |  |  |  |  |
| 288 | 9403_4426 | NA | NA | 35,319 | 35,829 |

bp, base pairs; NA, not applicable.

**S11 Table:** Read counts from amplicon sequencing detecting JCPyV early wraparound transcripts PML samples.

| Product length (bp) | Introns | ID (IMR-32) | ID (293) | Read<br>count<br>t<br>(case<br>1) | Read<br>count<br>t<br>(case<br>2) | Read<br>count<br>t<br>(case<br>3) | Read<br>count<br>t<br>(case<br>4) | Read<br>count<br>t<br>(case<br>5) | Read<br>count<br>t<br>(case<br>6) | Read<br>count<br>t<br>(case<br>7) | Read<br>count<br>t<br>(case<br>8) |
| --- | --- | --- | --- | --- | --- | --- | --- | --- | --- | --- | --- |
| 1359 | 9403_4426 | IMR-E16 | NA | 2 | 0 | 0 | 0 | 0 | 0 | 1 | 0 |
|  | 14533_9556 |  |  |  |  |  |  |  |  |  |  |
|  | 19663_14686 |  |  |  |  |  |  |  |  |  |  |
|  | 24793_19816 |  |  |  |  |  |  |  |  |  |  |
|  | 29923_24946 |  |  |  |  |  |  |  |  |  |  |
|  | 35053_30076 |  |  |  |  |  |  |  |  |  |  |
|  | 40183_35206 |  |  |  |  |  |  |  |  |  |  |
|  | 45313_40336 |  |  |  |  |  |  |  |  |  |  |
| 1206 | 9403_4426 | NA | NA | 2 | 1 | 3 | 0 | 1 | 0 | 10 | 2 |
|  | 14533_9556 |  |  |  |  |  |  |  |  |  |  |
|  | 19663_14686 |  |  |  |  |  |  |  |  |  |  |
|  | 24793_19816 |  |  |  |  |  |  |  |  |  |  |
|  | 29923_24946 |  |  |  |  |  |  |  |  |  |  |
|  | 35053_30076 |  |  |  |  |  |  |  |  |  |  |
|  | 40183_35206 |  |  |  |  |  |  |  |  |  |  |
| 1053 | 9403_4426 | IMR-E14 | NA | 28 | 1 | 5 | 7 | 12 | 1 | 58 | 20 |
|  | 14533_9556 |  |  |  |  |  |  |  |  |  |  |
|  | 19663_14686 |  |  |  |  |  |  |  |  |  |  |
|  | 24793_19816 |  |  |  |  |  |  |  |  |  |  |
|  | 29923_24946 |  |  |  |  |  |  |  |  |  |  |
|  | 35053_30076 |  |  |  |  |  |  |  |  |  |  |
| 900 | 9403_4426 | NA | NA | 161 | 8 | 13 | 14 | 103 | 9 | 278 | 96 |
|  | 14533_9556 |  |  |  |  |  |  |  |  |  |  |
|  | 19663_14686 |  |  |  |  |  |  |  |  |  |  |
|  | 24793_19816 |  |  |  |  |  |  |  |  |  |  |
|  | 29923_24946 |  |  |  |  |  |  |  |  |  |  |
| 747 | 9403_4426 | NA | NA | 997 | 55 | 47 | 64 | 529 | 37 | 916 | 733 |
|  | 14533_9556 |  |  |  |  |  |  |  |  |  |  |
|  | 19663_14686 |  |  |  |  |  |  |  |  |  |  |
|  | 24793_19816 |  |  |  |  |  |  |  |  |  |  |
| 594 | 9403_4426 | IMR-E15 | 293-E17 | 4,425 | 266 | 333 | 198 | 2,520 | 200 | 2,354 | 3,929 |
|  | 14533_9556 |  |  |  |  |  |  |  |  |  |  |
|  | 19663_14686 |  |  |  |  |  |  |  |  |  |  |
| 441 | 9403_4426 | NA | 293-E16 | 11,679 | 3,378 | 4,157 | 1,076 | 7,655 | 2,761 | 3,646 | 13,457 |
|  | 14533_9556 |  |  |  |  |  |  |  |  |  |  |
| 288 | 9403_4426 | NA | NA | 16,242 | 49,100 | 41,407 | 26,180 | 11,957 | 44,787 | 2,969 | 23,223 |

bp, base pairs; NA, not applicable.

**S12 Table:**
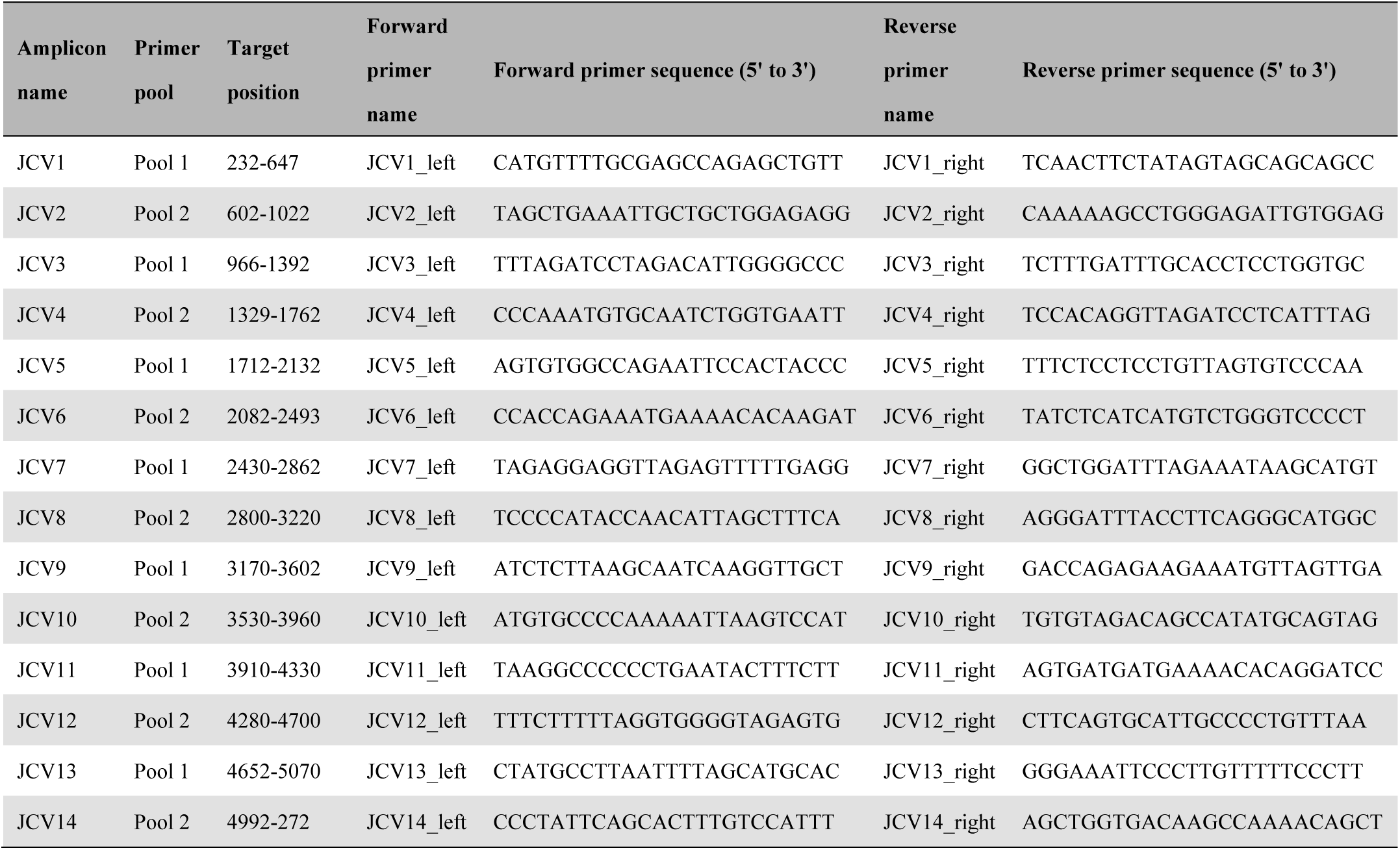
PCR primers used for amplicon sequencing of the JCPyV genome.

**S1 Fig:**
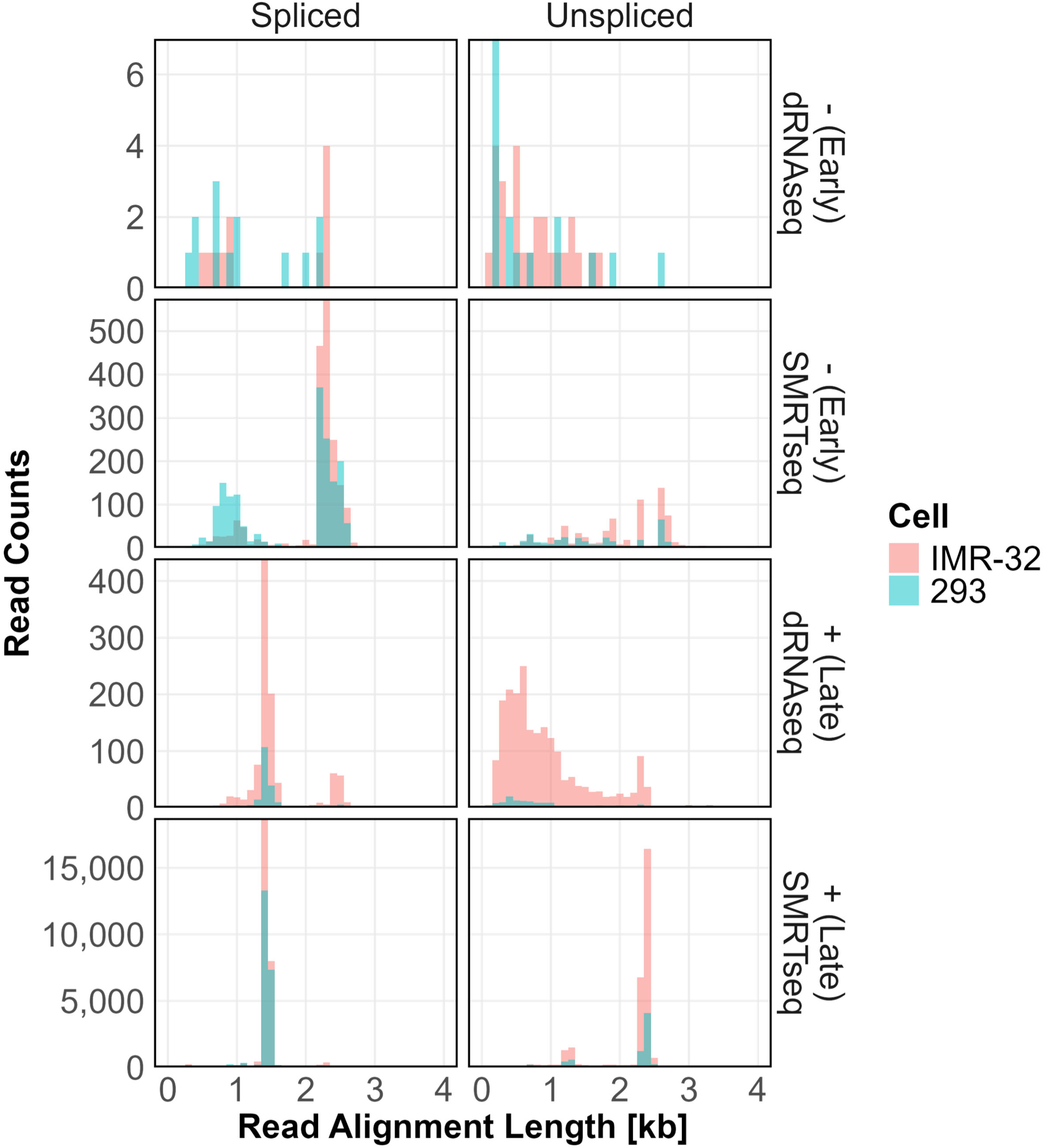
The distribution of read lengths aligned to the JCPyV genome. The distribution of mapped read lengths from IMR-32 (red) and 293 (green) cells are demonstrated based on the classification of the JCPyV genome strand (early or late), splicing status (spliced or unspliced), and sequencing technologies (dRNAseq or SMRTseq). The X-axis indicates the length of sequencing reads aligned to the JCPyV genome, whereas the Y-axis indicates the read counts of transcripts. dRNAseq, direct RNA sequencing; SMRTseq, single-molecule real-time sequencing.

**S2 Fig:**
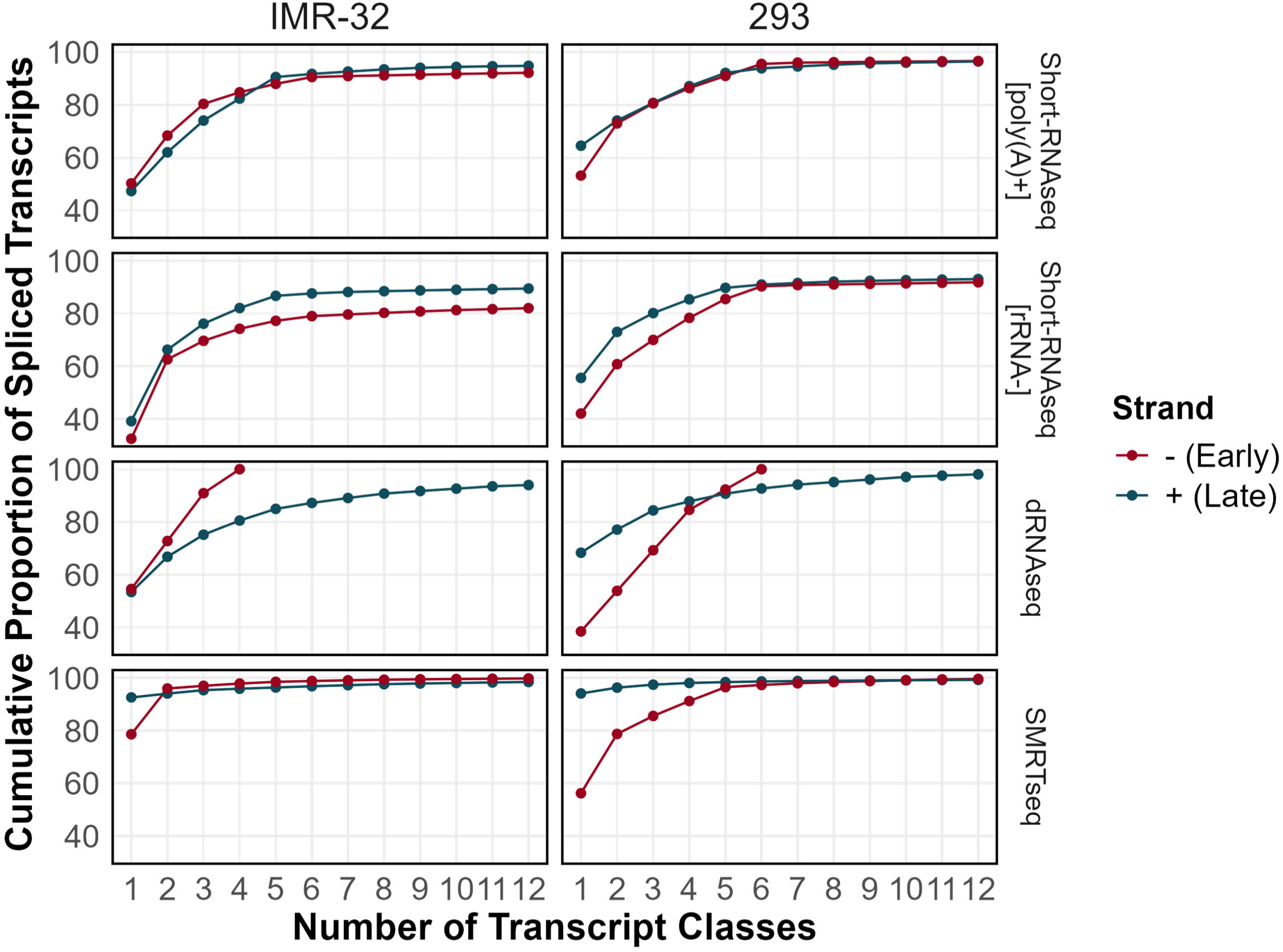
Cumulative proportions of viral transcripts. Cumulative proportions of spliced transcripts determined by each NGS method. Proportions were calculated for each strand. Red lines represent the negative strand (early region), while green lines represent the positive strand (late region). The results from IMR-32 cells (left panels) and 293 cells (right panels) are shown. Short-RNAseq, short-read RNA sequencing; rRNA, ribosomal RNA; dRNAseq, direct RNA sequencing; SMRTseq, single-molecule real-time sequencing.

**S3 Fig:**
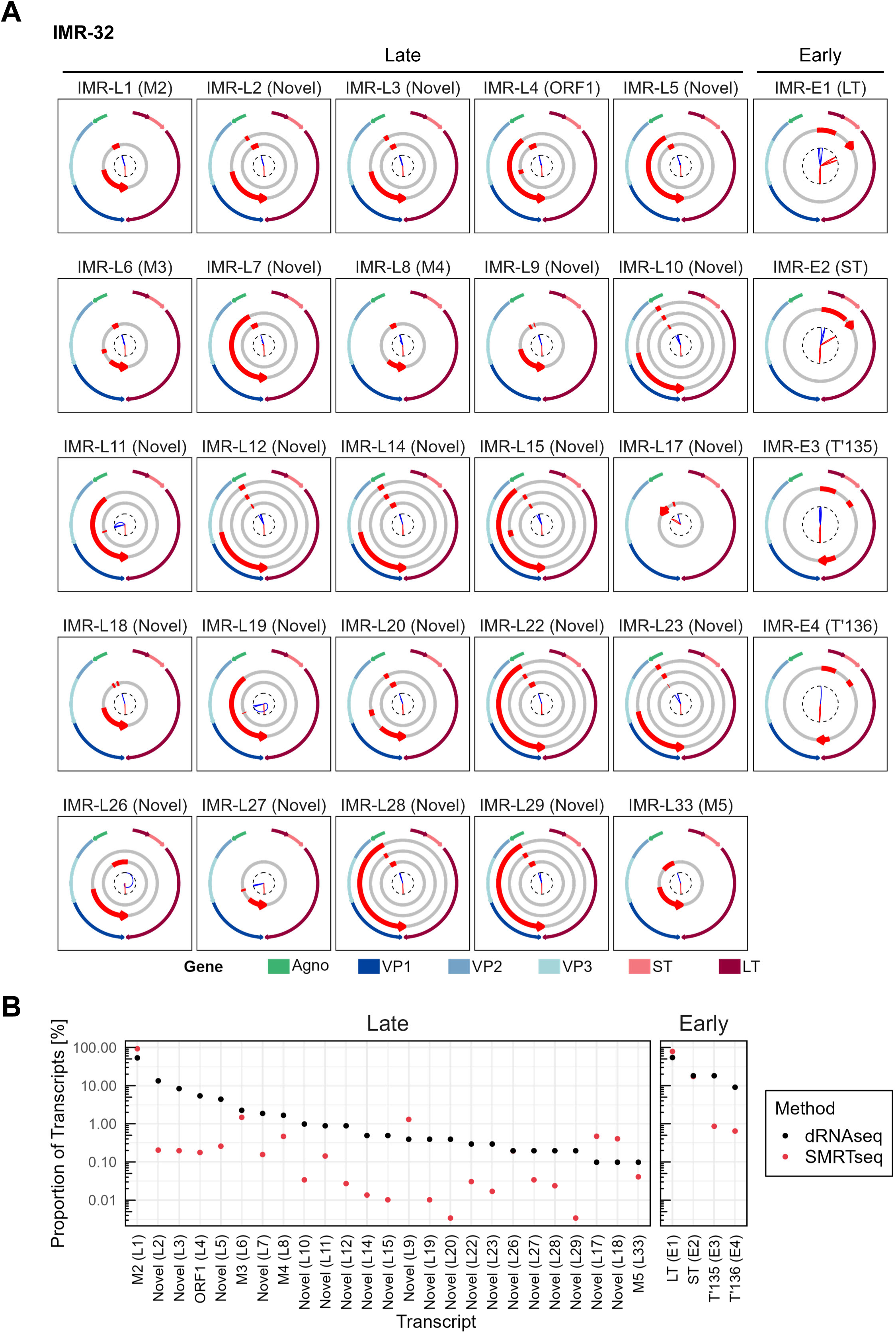
JCPyV transcript atlas in IMR-32 cells. **A.** Watch plots of viral transcripts detected by both dRNAseq and SMRTseq in IMR-32 cells, with each panel detailing each transcript class. Panels are presented in a similar manner to that described in **Fig 4A**. **B.** Relative abundance of viral transcripts demonstrated in **S3A Fig**. dRNAseq, direct RNA sequencing; SMRTseq, single-molecule real-time sequencing.

**S4 Fig:**
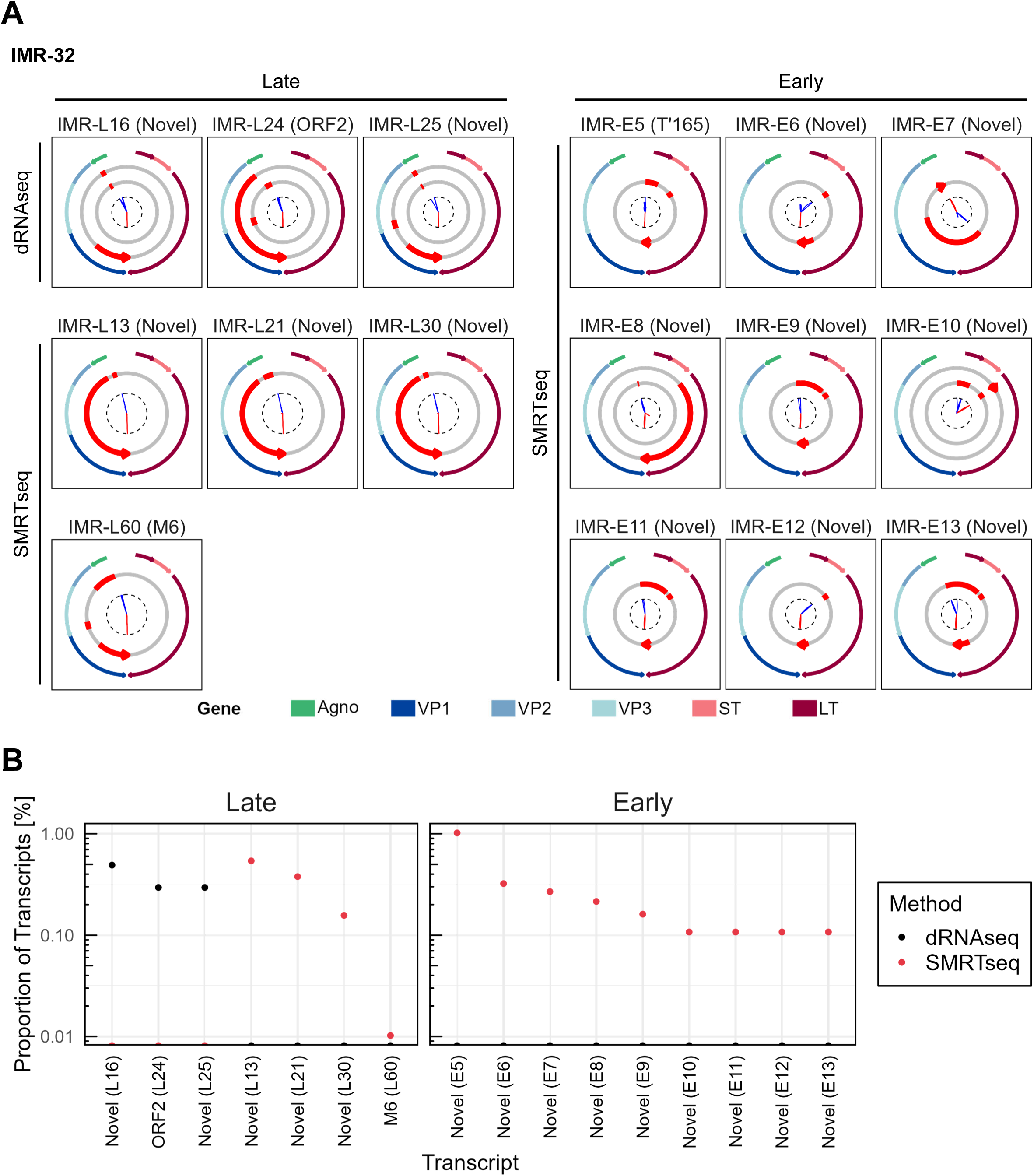
JCPyV transcripts detected only by dRNAseq or SMRTseq in IMR-32 cells. **A.** Watch plots of viral transcripts of JCPyV displayed in the same manner as described in **Fig 4A**. Late transcripts detected by dRNAseq (left upper), late transcripts detected by SMRTseq (left lower), and early transcripts detected by SMRTseq (right) are shown. **B.** Relative abundance of viral transcripts demonstrated in **S4A Fig**. dRNAseq, direct RNA sequencing; SMRTseq, single-molecule real-time sequencing.

**S5 Fig:**
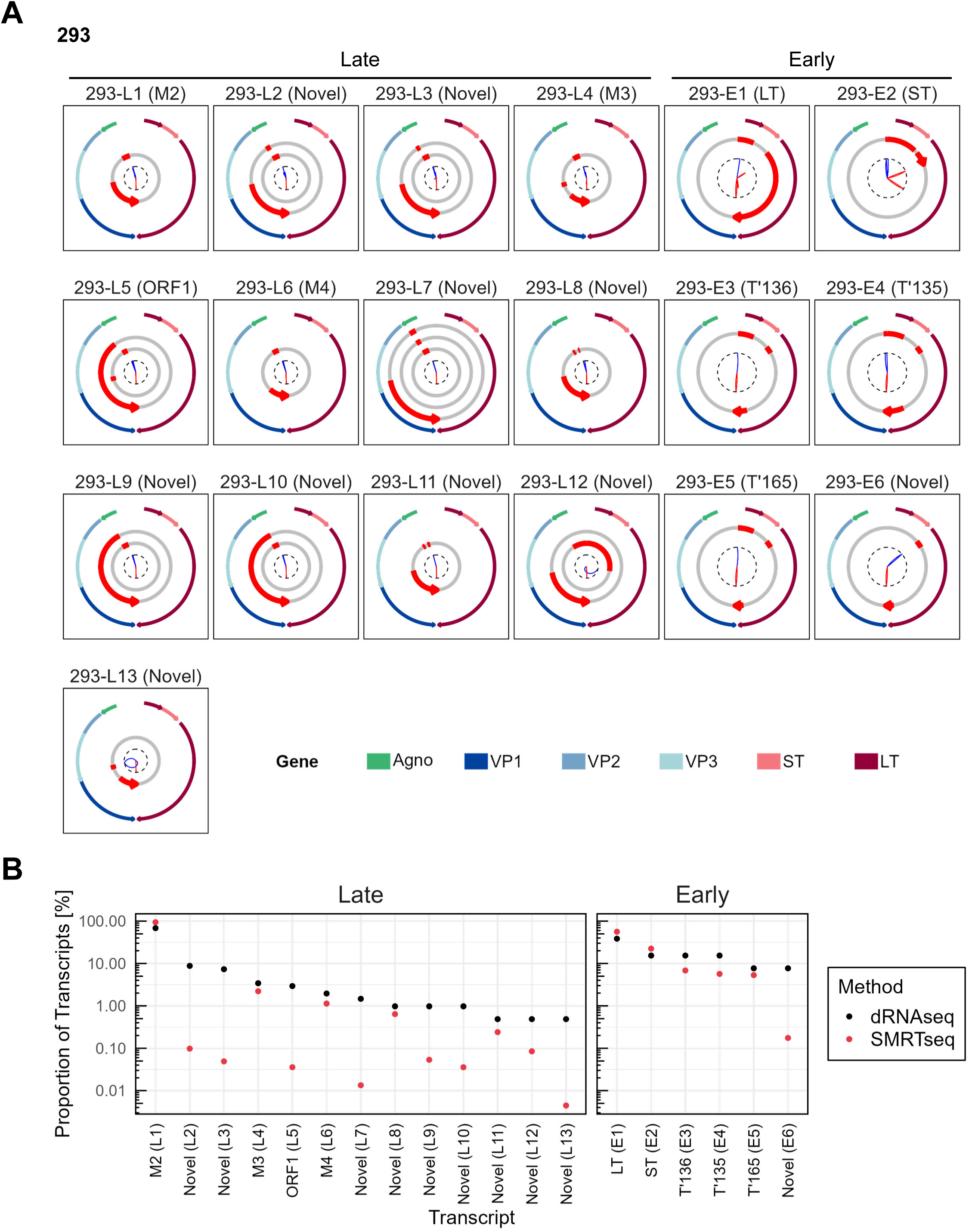
JCPyV transcript atlas in 293 cells. **A.** Watch plots of viral transcripts detected by both dRNAseq and SMRTseq in 293 cells, with each panel detailing each transcript class. Panels are presented in a similar manner to that described in **Fig 4A**. **B.** Relative abundance of viral transcripts demonstrated in **S5A Fig**. dRNAseq, direct RNA sequencing; SMRTseq, single-molecule real-time sequencing.

**S6 Fig:**
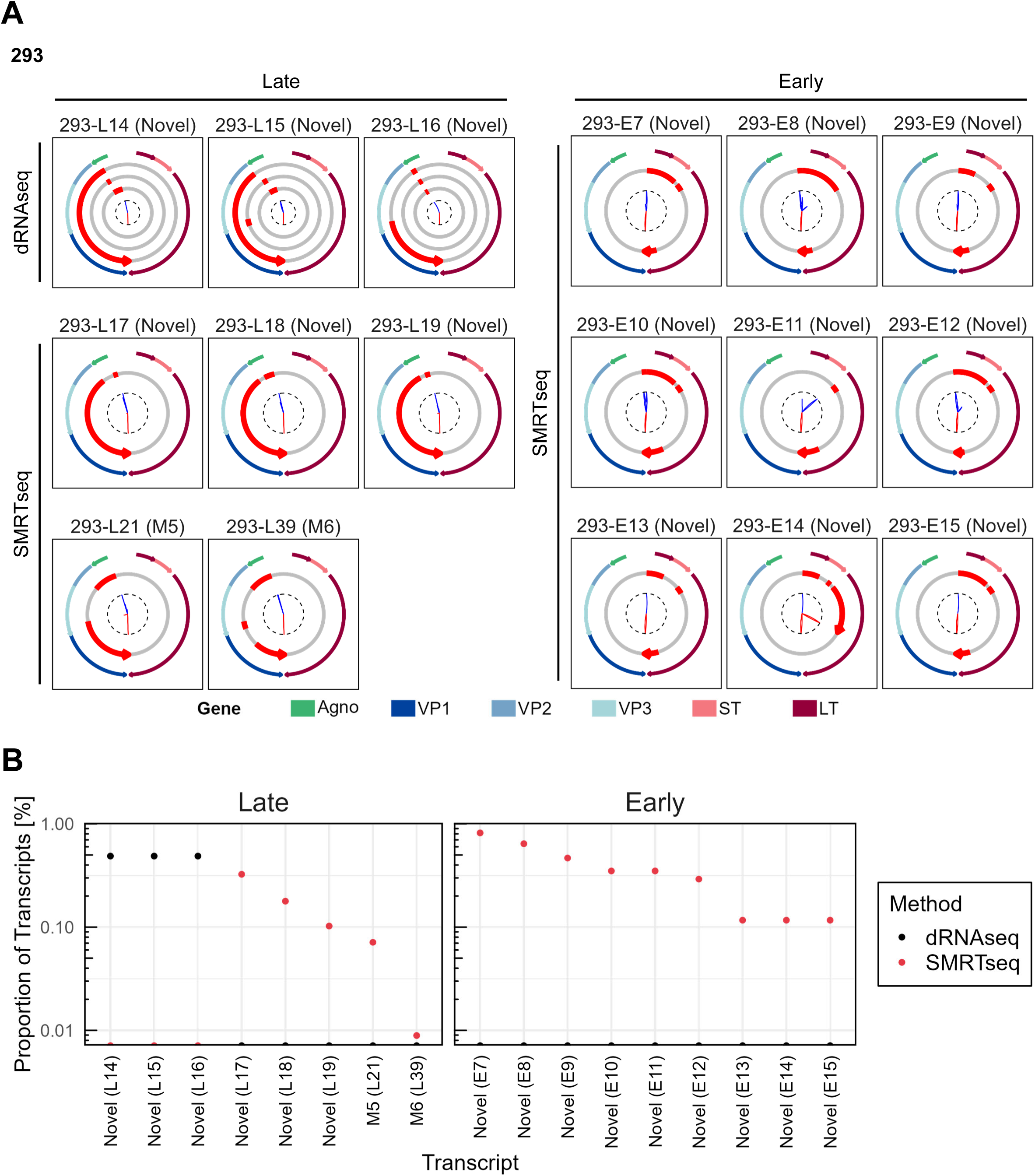
JCPyV transcripts detected only by dRNAseq or SMRTseq in 293 cells. **A.** Watch plots of viral transcripts of JCPyV displayed in the same manner as described in **Fig 4A**. Late transcripts detected by dRNAseq (left upper), late transcripts detected by SMRTseq (left lower), and early transcripts detected by SMRTseq (right) are shown. **B.** Relative abundance of viral transcripts demonstrated in **S6A Fig**. dRNAseq, direct RNA sequencing; SMRTseq, single-molecule real-time sequencing.

**S7 Fig:**
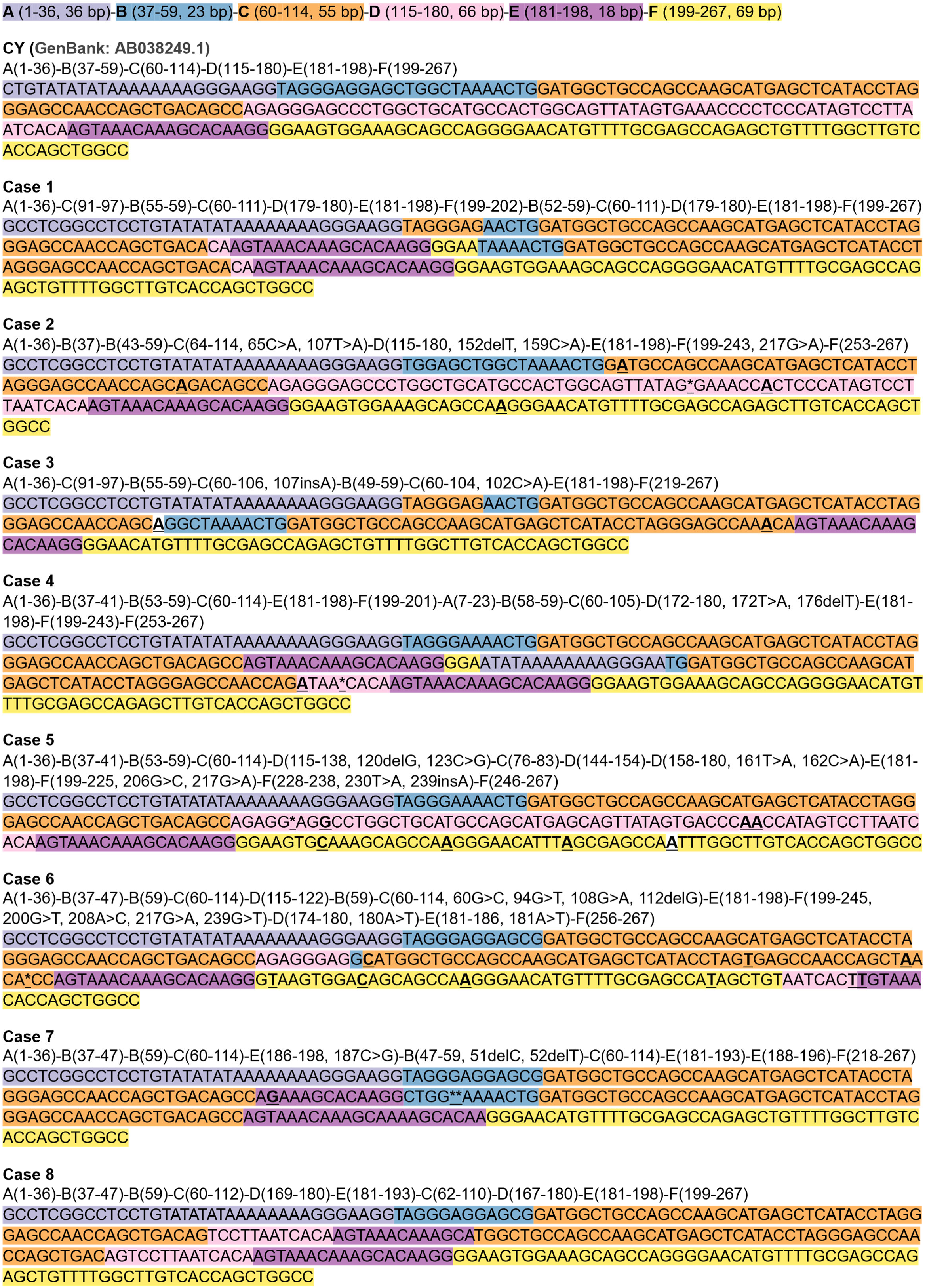
DNA sequence of the NCCR of the JCPyV genome in PML cases. DNA sequence of the NCCR, determined by amplicon sequencing, is shown. Each sequence block is highlighted in distinct colors. Nucleotide substitutions and insertions are shown in bold, while deletions are indicated by asterisks. Each number indicates the position in the JCPyV CY strain genome. The NCCR of the archetype (CY) JCPyV is also shown.

